# Cholinergic and noradrenergic modulation of perceptual decision making and learning

**DOI:** 10.64898/2026.06.22.732890

**Authors:** Ana Antonia Dias Maile, Evelina Andronova, Felix Ball, Dzhamilia Aravitska, Lisa K Dannenberg, Alfons Schnitzler, Jill X O’Reilly, Gerhard Jocham

**Author notes:** **Correspondence**: Ana Antonia Dias Maile.

## Abstract

Decision making often requires the integration of noisy bottom-up sensory evidence with top-down prior expectations learned from past experiences. Acetylcholine and noradrenaline have been proposed to shape these computations, yet it remains unclear whether they directly influence the weighting of sensory evidence and prior expectations during decision making or instead shape the learning process that gives rise to these expectations. To disentangle this, healthy participants (*n* = 62) completed a perceptual decision-making task under the influence of either the muscarinic acetylcholine receptor antagonist biperiden, the β-adrenoceptor antagonist propranolol, or placebo, in a within-subject crossover design. The task required the integration of sensory evidence with learned expectations. The reliability of these two sources of information varied independently. We show that both drugs led to a faster updating of current beliefs in response to new evidence. Under propranolol, these effects were specific to reward outcomes, whereas under biperiden, faster updating in response to both reward and non-reward outcomes resulted in less stable beliefs. Notably, neither drug affected the degree to which choices were governed by the strength of the sensory evidence. Together, these findings suggest that acetylcholine and noradrenaline guide perceptual decision making by modulating the updating of top-down prior expectations in response to new bottom-up evidence during learning.

## Introduction

Perception often relies on noisy and uncertain sensory input. For example, when someone waves at you in foggy weather, limited visibility may make it difficult to identify the person. However, past experience allows learning that certain people are more likely to be encountered in particular contexts. Being on a university campus, for instance, increases the expectation that the person is a fellow student or colleague. Perception and decision making therefore depend on the continuous integration of bottom-up sensory evidence with top-down prior expectations learned from past experiences.

Given the importance of this integration, previous research has been investigating its underlying neurochemical mechanisms. In particular, acetylcholine and noradrenaline have been proposed to play a key role in learning from past outcomes and, consequently, in the formation of top-down expectations. Consistent with this, muscarinic acetylcholine receptor antagonist has been shown to affect reward-guided decision making only when learning is required. Specifically, muscarinic receptor blockade reduces the stability of learned beliefs by accelerating their updating in volatile environments (Kurtenbach et al., 2026). Muscarinic acetylcholine receptor antagonist has further been shown to impair error-related behavioral and cortical adjustments (Danielmeier et al., 2015). Correspondingly, cholinergic responses in the basal forebrain have been shown to provide a precise reinforcement signal that scales with outcome expectancy (Hangya et al., 2015). Noradrenaline has been proposed to play an opposite role to this. Tonic noradrenergic activity facilitates attention-shifting to contextual changes, thereby promoting exploration and responses to new task-relevant stimuli (Aston-Jones & Cohen, 2005; Devauges & Sara, 1990). Additionally, a large body of evidence shows that noradrenaline plays a key role in synaptic plasticity, learning and memory consolidation (Jami et al., 2023; Larsen et al., 2023; Sara et al., 1999). In line with this, β-adrenoceptor antagonism has been shown to slow bottom-up updating of reward-contingencies in volatile environments (Lawson et al., 2021).

An alternative account proposes that acetylcholine and noradrenaline influence decision making by modulating how top-down expectations are integrated with bottom-up sensory evidence (Yu & Dayan, 2005). Within this framework, the effects of noradrenaline and acetylcholine on learning may arise from a differential weighting of the reliability of new bottom-up information relative to that of the current top-down belief. Both neuromodulators have been shown to play an important role in sharpening sensory neuronal representations (Eggermann et al., 2014; Hasselmo et al., 1997; Herrero et al., 2017; Kossl & Vater, 1989). Consequently, noradrenaline and acetylcholine would determine the extent to which incoming evidence updates current expectations. However, it remains unclear whether this integration is primarily linked to learning. Alternatively, acetylcholine and noradrenaline may govern the weighting of bottom-up information and top-down knowledge independently of learning at the time of decision making.

To disentangle pharmacological effects on evidence weighting from learning itself, we implemented a perceptual decision-making paradigm in which the reliability of bottom-up sensory evidence and top-down learned expectations within each decision was manipulated orthogonally. Because the weighting of each information source during decision making should depend on its reliability (de Lange et al., 2018; Hanks et al., 2011), this design allowed us to quantify their independent contributions to choice behavior. Additionally, learning of the top-down prior was independent of the strength of sensory evidence, as veridical feedback was given after each decision. This allowed us to measure effects on learning independently of perception and its integration. Participants completed the task across three pharmacological sessions in which they received the muscarinic acetylcholine receptor antagonist biperiden and the β-adrenoceptor antagonist propranolol in a randomized within-subject, placebo-controlled design. We investigated whether the drugs directly altered the weighting of sensory evidence and prior expectations during decision making or instead influenced reward-based learning. We expected biperiden to increase the learning rate causing a faster updating of prior beliefs, whereas propranolol was expected to reduce the learning rate leading to slower updating. Additionally, previous research has implicated both noradrenaline and acetylcholine in regulating spontaneous activity and evoked responses in sensory cortices. We therefore expected both drugs to reduce participants’ perceptual discrimination performance (Eggermann et al., 2014; Hasselmo et al., 1997; Herrero et al., 2017; Jimenez-Martin et al., 2021; Kasamatsu & Heggelund, 1982; Kossl & Vater, 1989; Kuchibhotla et al., 2017; McLean & Waterhouse, 1994; Meir et al., 2018; Waterhouse et al., 1998).

In brief, we find that both biperiden and propranolol increased learning rates, resulting in faster updating of prior beliefs. This accelerated updating was specific to reward outcomes under propranolol, whereas under biperiden it led to a maladaptive adjustment of prior beliefs across both reward and non-reward outcomes. Notably, neither drug affected the sensitivity to sensory evidence nor the integration of sensory evidence and prior expectations independently of learning at the time of choice.

## Results

To investigate the distinct contributions of acetylcholine and noradrenaline in perception, learning and their integration into a decision, 62 healthy participants performed a modified version of the random dot motion task (RDM). Participants were asked to indicate, on each trial, the mean movement direction of a noisy display of moving dots. We manipulated the quality of the sensory evidence by varying motion coherence (from 0 to 48%), the proportion of dots moving coherently to the left or right (Figure 1b).

**Figure 1:**
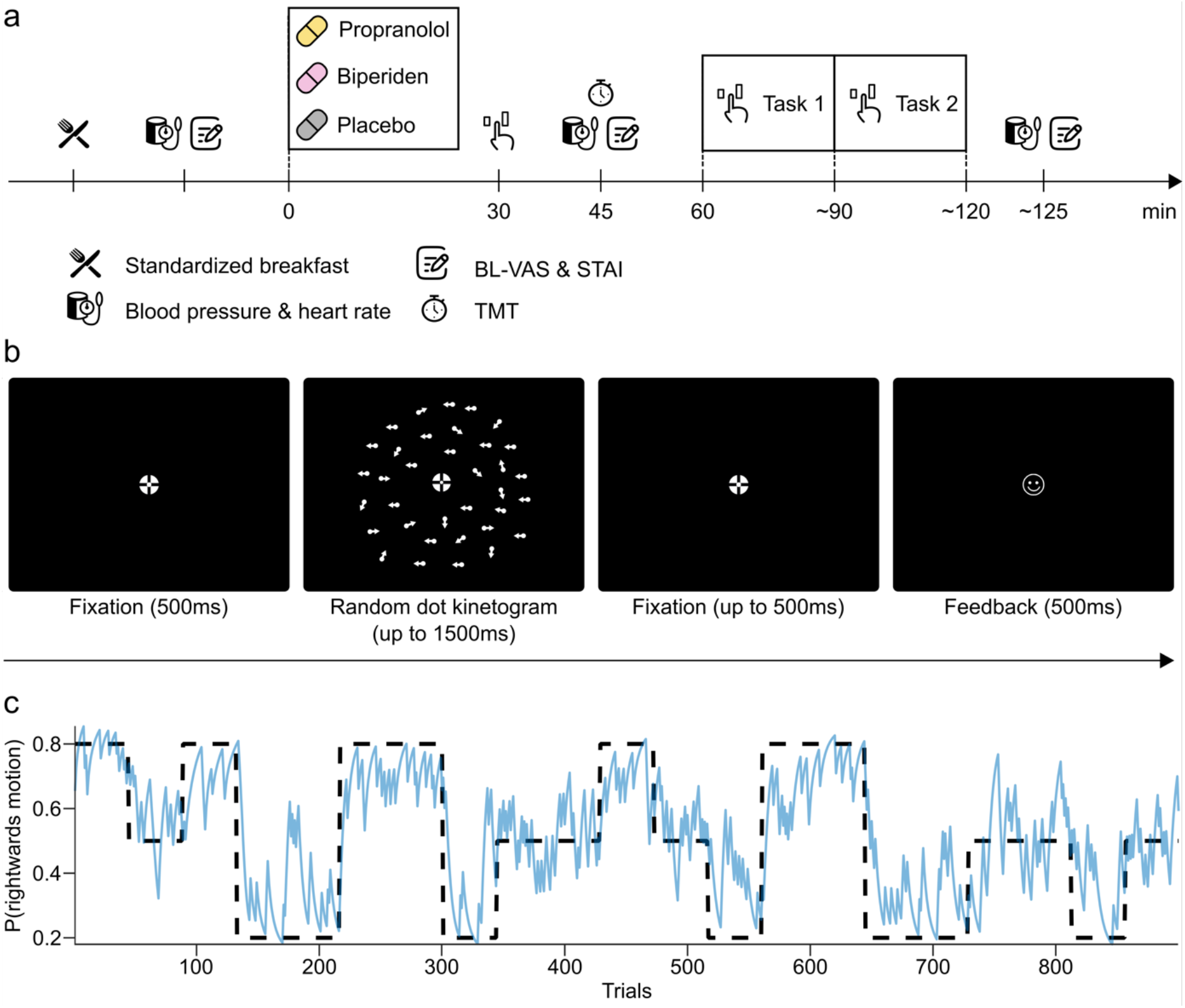
Study procedure and experimental task. **(a)** Timeline of experimental procedures (kept constant across sessions). Each testing day began with a standardized breakfast. Participants ingested the pill 1 h before performing the experimental tasks. Task order was constant within participants but counterbalanced across participants. Heart rate and blood pressure were monitored at three different time points, during which participants also completed the Bond & Lader visual analogue scales (BL-VAS) and the State-Trait Anxiety Inventory (STAI). Immediately prior to the first task, participants additionally performed the Trail-Making Test (TMT). These measures were used for control analyses (Supplement 2) **(b)** In the random dot motion task, participants were asked to indicate the mean movement direction of the random dot kinematogram. For this, they could use two sources of information: the coherent motion and a prior probability of a left- or rightward movement, which could be inferred based on the outcomes of previous trials. Participants were allowed to respond until up to 500 ms after the offset of the kinematogram (2000 ms response deadline altogether) **(c)** Example of the time-varying prior probability of movement direction, which switched every 44 or 84 trials. The dashed line represents the true prior and the solid blue line is the estimated prior from a Bayesian optimal learner.

In addition, the task consisted of periods in which both movement directions were equally likely and periods in which either a left- or rightward motion was more probable (prior probability, Figure 1c). Changes between these periods were not signaled to participants but needed to be learned. Therefore, decisions could be based on both the available sensory evidence and the learned prior expectation (see Figure 1b-c; similar to Froböse & Ort, 2022; Marshall et al., 2024). We reasoned that statistical knowledge based on prior experience would be particularly beneficial on trials with low strength of sensory evidence. Participants performed the task on three separate sessions, where they received either a single oral dose of the M_1_-preferring muscarinic receptor antagonist biperiden (4mg), the β-adrenoceptor antagonist propranolol (40mg), or a placebo in a randomized, within-subject design (Figure 1a).

### Choice behavior is affected both by motion coherence and prior expectations

We first confirmed that participants’ decisions were indeed consistently influenced by both the level of motion coherence and the learned prior belief. A shift in decision making to favor the motion direction of the prior was evident in the raw task behavior (Figure 2a). This was confirmed using a general linear mixed-effects model (GLMM) investigating the influence of motion coherence and prior on choice (left vs. right) across drug conditions. Since the prior was not explicitly signaled to participants, here, we estimated it using a Bayesian optimal learner, which captured what participants could ideally know. Participants’ estimates of prior left- or rightward motion probability may well deviate from these normative estimates, a point that we will investigate in further detail in subsequent sections. Model results corroborated that coherence level and the estimated prior had a significant main effect on choice behavior (coherence level: β = 2.08, *SEM* = 0.11, *z*(166673) = 18.42, *p* < 0.001; prior: β = 0.59, *SEM* = 0.04, *z*(166673) = 16.19, *p* < 0.001; marginal *R^2^_GLMM_* = 0.593; Figure 2c and Table S1). With increasing coherence and prior strength, participants were more likely to choose the corresponding motion direction (Figure 2a). Furthermore, participants appeared to put more weight on the perceptual information (i.e., the coherent motion) as on the learned prior, as indicated by the higher regression weight (Figure 2c).

**Figure 2:**
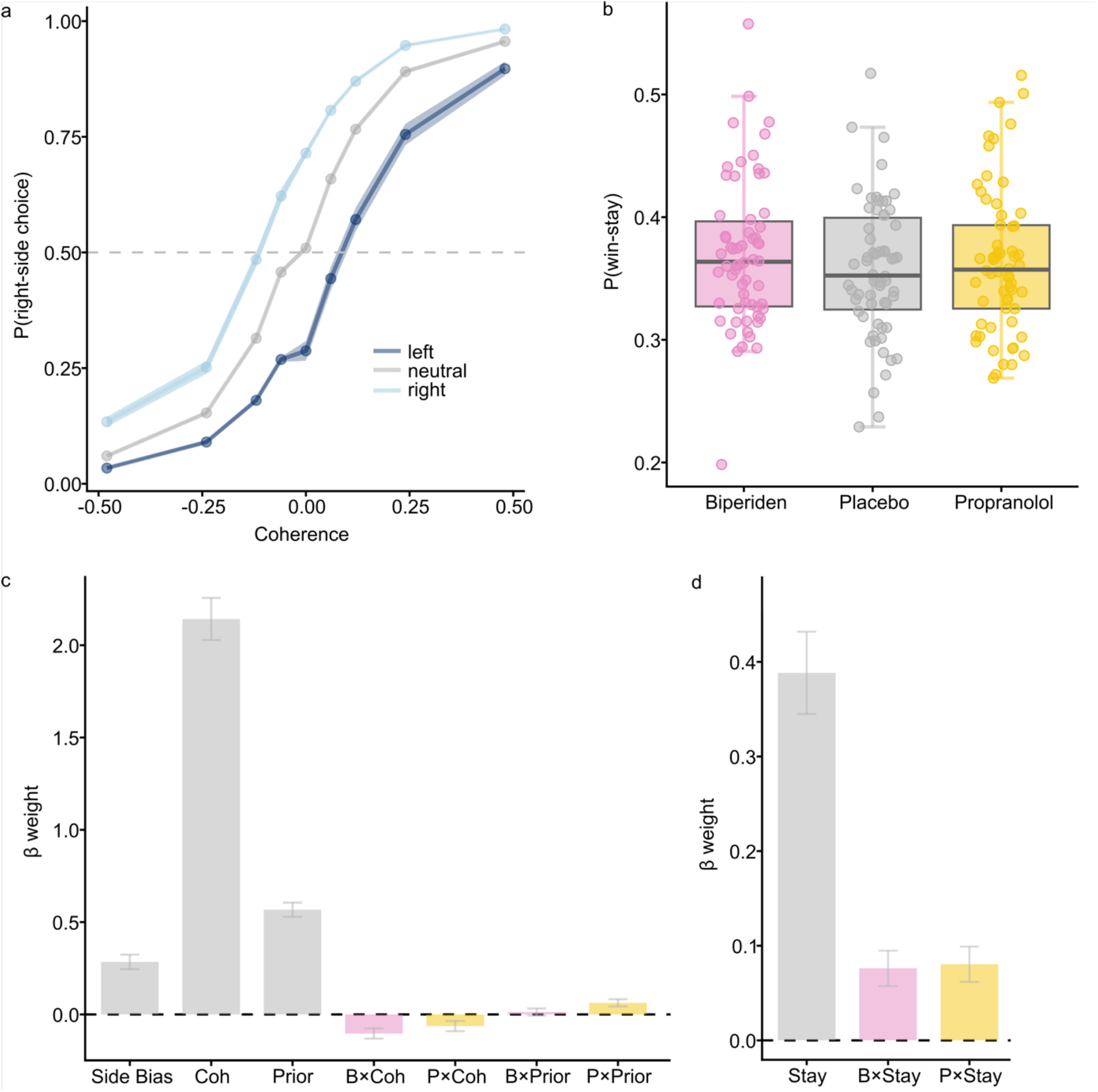
Drug effects on choice behavior. **(a)** Raw task behavior, averaged across drug condition: Probability for a right-side choice as a function of motion coherence and prior probability (shades of blue, Bayes-optimal prior favoring a left or right choice, or neutral prior where both directions are equally likely on average). Negative coherence levels correspond to leftward motion. Shades depict the standard error. **(b)** Raw prior influence, averaged across drug condition: Difference in the probability of a correct choice between trials in which Bayes-optimal prior was aligned with versus opposed to motion coherence. Light blue lines represent individual participant. The thick blue line indicates the group mean and the shaded area reflects the standard error. **(c)** Raw win-stay behavior, separated by drug condition. The probability of a win-stay choice for each drug condition, with individual participants represented as dots and distributions summarized by boxplots. Win-stay behavior was increased under both propranolol and biperiden relative to placebo. **(d)** GLMM of participants choice behavior (right-side choices). Bars represent regression coefficients (β) with standard errors. Choices are influenced by both motion coherence (Coh) and prior belief (Prior), with a stronger effect of motion coherence. The effect of coherence on choice behavior was reduced under propranolol (P) and biperiden (B) relative to placebo, as indicated by the significant negative interaction terms. In contrast, the influence of prior was increased under propranolol, as reflected by a significant positive interaction. **(e)** GLMM of win-stay behavior. Bars represent regression coefficients (β) with standard errors. Following reward outcomes, participants tended to repeat the same choice (stay), which was further increased under propranolol (P) and biperiden (B) as indicated by the significant positive interaction terms.

Furthermore, participants flexibly integrated the coherent motion and the learned prior when making decisions. Choice behavior (Figure 2a) clearly indicated that, under lower levels of perceptual reliability (i.e., at low motion coherence), participants relied more on the learned prior probability. To quantify this relationship, we set up a GLMM in which we asked whether the increase in the probability of making a correct choice as a function of (unsigned) motion coherence differed between congruent and incongruent trials (i.e., trials in which the prior indicated the same vs. the opposite choice as the direction of coherent motion). This analysis confirmed that the difference in the percentage of correct choices between congruent and incongruent trials decreased with increasing motion coherence (β = -0.66, *SEM* = 0.05, *t*(185) = -13.73, *p* < 0.001; marginal *R^2^_GLMM_* = 0.235; Figure 2a-b and Table S2).

### Biperiden and propranolol both appear to affect the balance between sensory evidence and prior belief

Turning to interactions between coherence, prior and drug condition in the same GLMM, we found that both biperiden and propranolol reduced the degree to which choices were governed by sensory evidence (effect of coherent motion, β = -0.10, *SEM* = 0.03, *z*(166658) = -3.76, *p* < 0.001; β = -0.01, *SEM* = 0.03, *z*(166658) = -2.26, *p* = 0.024; for the interaction of biperiden and propranolol with coherence level, respectively, marginal *R^2^_GLMM_* = 0.594; Figure 2c and Table S3). This seems to suggest that both drugs reduce participants’ sensitivity to the strength of the sensory evidence. However, we will show below that this apparent effect on perceptual sensitivity may instead reflect an underlying effect on learning. In addition, propranolol increased the effect of the learned prior on choices (β = 0.06, *SEM* = 0.02, *z*(166658) = 3.23, *p* = 0.001; marginal *R^2^_GLMM_* = 0.594; Figure 2c and Table S3). This may indicate a stronger influence of learned prior expectations on participants’ choices under propranolol, in line with our predictions.

### Biperiden and propranolol may affect learning, rather than weighting, of the prior

Previous work showed that biperiden affects reward-guided decision making by modifying the learning rate, rather than affecting decisions themselves (Kurtenbach et al., 2026). Similarly, noradrenergic transmission is involved in learning under conditions of changing contingencies (Glennon et al., 2019; Jepma et al., 2016) Therefore, an alternative explanation for the apparent downweighting of sensory evidence compared to the prior (Figure 2) is that the prior is inaccurately learned in the drug condition. The estimates of prior probability used above were based on a Bayesian optimal learner, which captured what participants could ideally know (Behrens et al., 2007). Hence, this drug effect on prior probability may either reflect a change in the use of this information or a change in learning i.e., how closely participants’ estimates of prior probability aligned with the Bayesian optimal learner.

We therefore used, in a first step, a model-agnostic approach to determine whether the degree to which past outcomes were used to guide future choices was affected by drug condition, namely a win-stay/lose-switch analysis. Faster updating of the prior belief is reflected in an increased likelihood of repeating a previously rewarded choice on the next trial (win-stay), whereas previously non-rewarded choices are related to a higher probability of switching (lose-switch; Estes, 1957). Conversely, slower updating is characterized by reduced win-stay/lose-switch behavior. Drug effects were analyzed separately for trials following rewarded and non-rewarded outcomes to assess potential differences. We observe increased win-stay behavior under both biperiden and propranolol (β = 0.09, *SEM* = 0.02, *z*(133447) = 4.07, *p* < 0.001; β = 0.08, *SEM* = 0.02, *z*(133447) = 4.30, *p* < 0.001; for the interaction of biperiden and propranolol with stay bias, respectively, marginal *R^2^_GLMM_* = 0.622; Figure 2b,d and Table S4). This pattern suggested faster updating of prior beliefs in response to reward outcomes. While this finding was consistent with the hypothesis for biperiden, it contradicted the prediction for propranolol where we expected a slower updating of the prior. In contrast, neither drug significantly modulated lose-switch behavior (β = - 0.04, *SEM* = 0.04, *z*(32971) = -1.02, *p* = 0.309; β = 0.02, *SEM* = 0.04, *z*(32971) = 0.49, *p* = 0.627; for the interaction of biperiden and propranolol with stay bias, respectively, marginal *R^2^_GLMM_* = 0.568; Table S5). The effect therefore did not extend to non-reward outcomes, where a faster updating would be expected to reduce stay behavior after errors.

Notably, the apparent drug modulation of motion coherence was reduced in this win-stay/lose-switch analysis, where we approximated learning by accounting for how past outcomes guide current choices, supporting the interpretation that the apparent effect of drug on weighting of the sensory evidence in fact reflects misestimation of the prior. Only biperiden significantly modulated the influence of motion coherence on choice behavior by decreasing the degree to which choices were governed by sensory evidence in response to reward outcomes (interaction of biperiden with coherence in trials following reward: β = -0.12, *SEM* = 0.03, *z*(133447) = -3.72, *p* < 0.001; for all other drug interactions with coherence following rewarded or unrewarded trials: β ≥ - 0.07, *SEM* ≥ 0.03, *z* ≥ -1.34, *p* ≥ 0.182).

Participants additionally exhibited a bias toward rightward motion, as indicated by a significant intercept across all models (Figure 2c, intercept from the choice model including all trials: β = 0.28, *SEM* = 0.04, *z*(166658) = 7.24, *p* < 0.001; marginal *R^2^_GLMM_* = 0.594; Figure 2c and Table S3). However, this bias was not modulated by either of the drugs (β = 0.03, *SEM* = 0.04, *z*(166658) = 0.69, *p* = 0.492; β = -0.03, *SEM* = 0.04, *z*(166658) = -0.71, *p* = 0.481; for the main effect of biperiden and propranolol in the choice model including all trials, respectively, marginal *R^2^_GLMM_* = 0.594). Nevertheless, given its pronounced influence on choice behavior, the side bias was taken into account in the subsequent computational modeling.

We additionally confirmed that drug effects were independent of the following potential confounds: All cholinergic and noradrenergic effects on participants’ choices were independent of drug effects on mood, state anxiety, visuomotor processes, heart rate, blood pressure and session order (see Supplement 2 for control analyses). Further, the drugs did not significantly influence participants’ reaction times (Table S6).

In sum, participants appear to update their prior expectations faster in response to rewards under both drugs compared to placebo. Therefore, both drugs seem to modulate learning, rather than weighting, of the prior.

### Computational learning model confirms that biperiden and propranolol affect learning rate rather than evidence weighting

While our win-stay analyses provided model-agnostic evidence for drug effects on learning, they only captured effects of the outcome from the immediately preceding trial. To fully account for how participants inferred beliefs about prior probability across multiple trials and how this was integrated with sensory evidence, we implemented computational models. This allowed us to determine whether the observed drug effects reflect changes in learning and/or in the flexible weighting of the two information sources (motion coherence and learned prior). The preregistered model implemented a Q-learning with a delta update rule governed by a learning rate (see methods, equation 1). Given the reward-specific effects of both drugs in the model-free win-stay analyses above, we additionally tested a model variant with separate learning rates for rewarded and unrewarded trials. However, a joint learning rate provided a better fit to the data (BIC = 730.39 and BIC = 732.04 for median Bayes Information Criterions (BIC) of the models with one learning or separate learning rates, respectively). Consistent with the flexible influence of the prior depending on perceptual reliability observed in participants’ behavior (Figure 2a-b), the model assumed an additive value construction. In particular, the evidence in favor of either a left or right choice was jointly governed by (signed) motion coherence and prior belief, with their relative contribution being determined by the weighting parameter θ (values closer to 0 indicated a stronger reliance on the prior and values closer to 1 indicated a stronger reliance on sensory evidence). Choices were generated using a softmax choice rule with an inverse temperature τ capturing choice stochasticity and a side bias parameter b capturing left- or right-choice biases.

The learning rate λ was significantly increased under both biperiden (*Mdn_biperiden_* = 0.38, *Mdn_placebo_* = 0.28, *z* = 3.14, *p* = 0.002, *r* = 0.40) and propranolol (*Mdn_propranolol_* = 0.33, *z* = 2.09, *p* = 0.037, *r* = 0.26; Figure 3a-b). This indicated that participants updated their prior estimate faster under both drugs, consistent with the win-stay/lose-switch analysis. To assess whether this pattern led to estimates of the prior that more closely tracked the optimal estimates, we computed the mean squared errors (MSE) of each participant’s trial-wise prior estimates and compared them between placebo and drugs. Under biperiden, the MSE was significantly higher than under placebo suggesting that the increased learning rate led to worse prior estimates, making it maladaptive (*Mdn_biperiden_* = 0.03, *Mdn_placebo_* = 0.02, *z* = 3.17, *p* = 0.002, *r* = 0.23; Figure 3c). In contrast, there was no significant difference in the MSE between propranolol and placebo (*Mdn_propranolol_* = 0.03, *z* = 1.52, *p* = 0.127, *r* = 0.11; Figure 3c).

**Figure 3:**
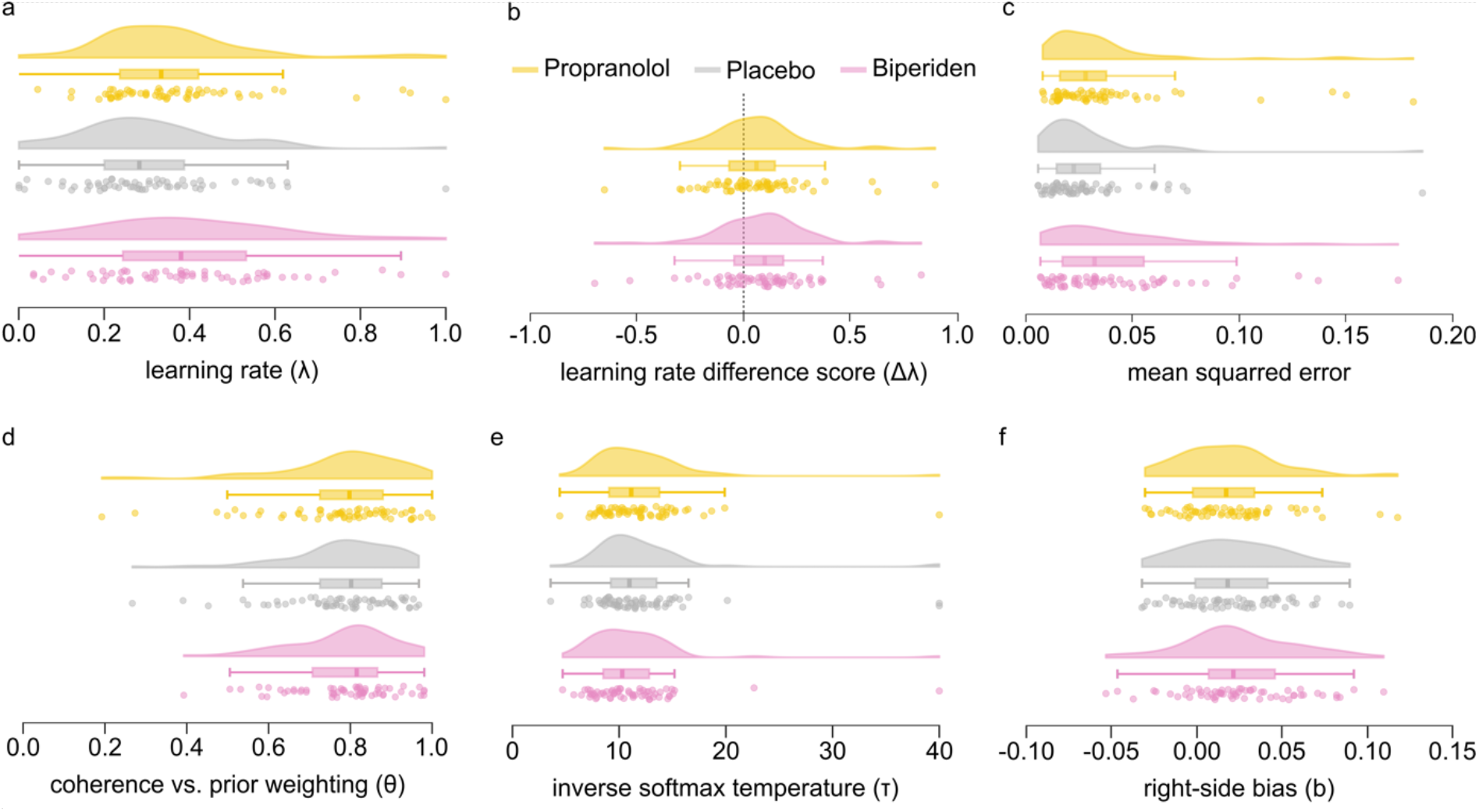
Parameter estimates per participant and drug conditions. Raincloud plots in **a**, **d**, **e**, **f** illustrate the distribution of the four model parameters across drug conditions: **(a)** learning rate (λ), **(d)** relative weighting (θ), **(e)** inverse softmax temperature (τ) and **(f)** right-side bias (b). Panel **b** shows difference scores for the learning rate (Δλ; drug-placebo), with the dotted line marking zero (no difference). Panel **c** displays the distribution of mean squared error (MSE) across drug conditions. Half-violins and boxplots summarize the distribution of single-subject values (dots) for each drug condition. **(a,b)** Learning rates were significantly higher under biperiden and propranolol compared to placebo. **(c)** This increased updating was maladaptive under biperiden, as reflected by significantly higher MSE relative to placebo. **(d)** The relative weighting of perceptual information and the prior beliefs was not significantly modulated by drug condition. **(e)** Choices tended to be more stochastic under biperiden, reflected in a trend for a lower inverse softmax temperature compared to placebo. There was no significant effect for propranolol. **(e)** Participants displayed a right-side choice bias that was not significantly modulated by drug condition.

### Differential drug effects on learning from positive and negative outcomes

Given that the effects we observed above in our model-free analyses were specific to win-stay behavior, we further examined the model with separate learning rates for rewarded and unrewarded choices. While this model did not improve the fit compared to the joint learning rate model, it allowed us to assess whether changes in learning were specific to reward or non-reward outcomes, or whether these changes occurred regardless of outcome type. This model revealed that propranolol specifically increased learning rates for rewarded choices (*Mdn_propranolol_* = 0.31, *Mdn_placebo_* = 0.30, *z* = 1.98, *p* = 0.048, *r* = 0.25; Figure 4a-b) with no significant effect on the unrewarded learning rate (*Mdn_propranolol_* = 0.28, *Mdn_placebo_* = 0.24, *z* = 1.07, *p* = 0.285, *r* = 0.14; Figure 4c-d). Biperiden, in contrast, increased learning rates for both rewarded (*Mdn_biperiden_* = 0.36, *z* = 2.53, *p* = 0.012, *r* = 0.32; Figure 4a-b) and unrewarded trials (*Mdn_biperiden_* = 0.38, *z* = 2.22, *p* = 0.027, *r* = 0.28; Figure 4c-d), with larger effects for rewarded trials (see Supplement 3 for complete model description).

**Figure 4:**
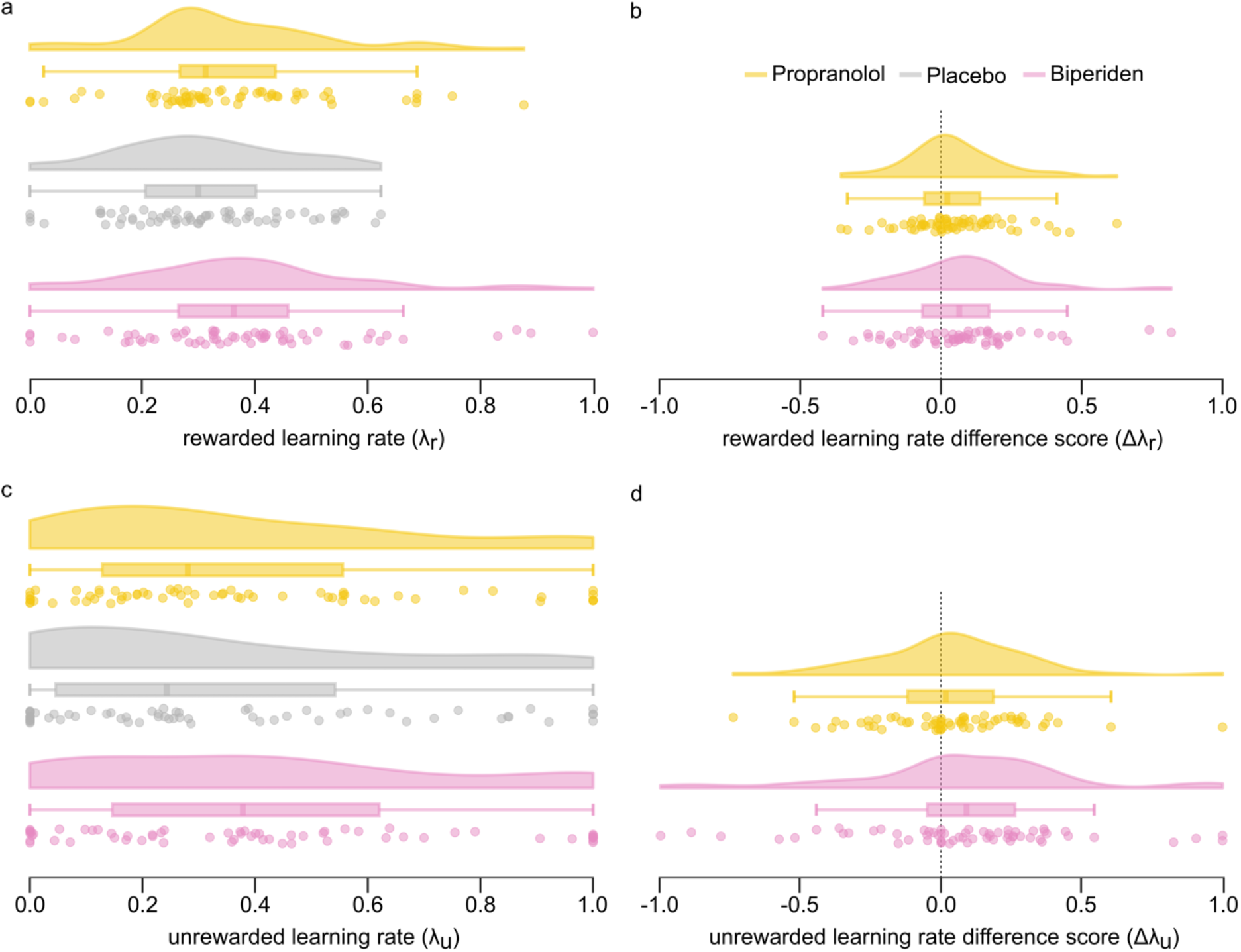
Different learning rates for reward and non-reward outcomes per participant and drug condition. Raincloud plots in **a** and **c** illustrate the distribution of rewarded **(a)** and unrewarded **(c)** learning rate across drug conditions. Panels **b** and **d** shows difference scores for the rewarded **(b)** and unrewarded **(d)** learning rate (Δλ_r_ and Δλ_u_; drug-placebo), with the dotted line marking zero (no difference). Half-violins and boxplots summarize the distribution of single-subject values (dots) for each drug condition. **(a,b)** Learning rates of rewarded trials were significantly higher under biperiden and propranolol compared to placebo. **(c,d)** Learning rates of unrewarded trials were significantly higher under biperiden, with no significant effect under propranolol.

### Once differences in learning rate are accounted for biperiden and propranolol do not affect the weighting of sensory and prior evidence

Based on the GLMM, we expected a relative shift in the weighting of coherence and prior under propranolol. However, the relative weighting was not significantly modulated by either propranolol (*Mdn_propranolol_* = 0.80, *z* = -0.87, *p* = 0.383, *r* = 0.11) or biperiden (*Mdn_biperiden_* = 0.82, *z* = -0.11, *p* = 0.914, *r* = 0.01; Figure 3d). Hence, there was no cholinergic or noradrenergic effect on the relative weight given to sensory evidence versus prior probability, contrary to our hypotheses. However, regardless of drug condition and consistent with the results from the GLMM, participants’ choices were influenced more strongly by coherent motion than by the learned prior, as indicated by values of the weighting parameter θ close to 1 (*Mdn_placebo_* = 0.80).

The inverse softmax temperature (τ), capturing choice stochasticity, was not significantly modulated by propranolol (*Mdn_propranolol_* = 10.29, *Mdn_placebo_* = 10.96, *z* = 0.14, *p* = 0.886, *r* = 0.02). There was a trend toward a lower inverse temperature and thus an increased choice stochasticity under biperiden (*Mdn_biperiden_* = 11.13, *z* = -1.84, *p* = 0.066, *r* = 0.23; Figure 3e). Participants additionally exhibited a rightward choice bias, which was not modulated by either drug (*Mdn_placebo_* = 0.02, biperiden: *Mdn_biperiden_* = 0.16, *z* = 1.54, *p* = 0.124, *r* = 0.20; propranolol: *Mdn_propranolol_* = 0.02, *z* = -0.47, *p* = 0.636, *r* = 0.06; Figure 3f), consistent with the GLMM.

Parameter recovery and model validation confirmed the reliability of the fitting procedure (Supplement 4). Simulated behavior successfully recovered the ground-truth parameters and reproduced key behavioral findings including the drug effects on motion coherence observed in the GLMM. Importantly, the drug modulation of coherence effects on choice observed in the GLMM could be fully explained by drug-dependent changes in the learning rate. This aligns with the win-stay/lose-switch analysis where drug modulations of coherence were also less robust. Together, this suggests that the observed drug-induced changes of coherence effects on choice we reported above using our regression-based approach do not reflect a genuine change in sensitivity to sensory inputs and instead reflects altered learning of environmental contingencies.

Taken together, propranolol and biperiden significantly altered learning dynamics, with both drugs causing a faster updating of prior beliefs. Notably, once this learning effects was accounted for, neither propranolol nor biperiden modulated the relative weighting of perceptual and learned information at the time of choice. Biperiden tended to additionally increase choice stochasticity.

## Discussion

Decision making relies on learning from past experiences and integrating this prior knowledge with current perceptual evidence. Noradrenaline and acetylcholine, which modulate both perception and learning, are thought to guide this integration (Eggermann et al., 2014; Hangya et al., 2015; Kossl & Vater, 1989; Lawson et al., 2021; McLean & Waterhouse, 1994; Meir et al., 2018; Yu & Dayan, 2005). In the present study, we investigated the role of these neurotransmitter systems in a perceptual decision-making task that required the integration of sensory evidence with prior expectations learned across past outcomes. Participants received the muscarinic acetylcholine receptor antagonist biperiden and the β-adrenoceptor antagonist propranolol in a within-subject, double-blind, placebo-controlled, crossover design. We investigated drug effects using two complementary approaches: First, GLMMs allowed us to examine how task parameters influenced behavior and how these effects were modulated by the pharmacological manipulations. Second, computational modeling allowed us to analyze drug effects on the underlying computations, including learning and the flexible integration of prior knowledge and sensory information. Our results highlight the role of noradrenaline and acetylcholine in learning and updating of top-down prior beliefs based on new bottom-up evidence. Specifically, muscarinic acetylcholine receptor antagonist under biperiden and β-antagonism under propranolol lead to a faster updating of prior beliefs as indicated by increased learning rates under both drugs compared to placebo. Therefore, biperiden and propranolol caused prior estimates to be influenced more strongly by recent outcomes compared to more distant experiences. Moreover, modulation of learning rates under biperiden led prior estimates to deviate more from the statistical optimum compared to placebo. Hence, increased learning rates under biperiden caused maladaptive adjustments in estimates of the prior. Prior estimates under propranolol did not significantly deviate relative to the statistical optimum compared to placebo possibly due a less pronounced effect on learning rates or to the selectivity for reward outcomes: Following a rewarded choice, prior estimates were adjusted faster under propranolol compared to placebo. Updates following non-rewarded choices however did not significantly differ between propranolol and placebo.

Perceptual discriminability has been shown to depend on cholinergic and noradrenergic modulation of the signal-to-noise-ratio in sensory cortices (Eggermann & Feldmeyer, 2009; Hasselmo et al., 1997; Kasamatsu & Heggelund, 1982; Meir et al., 2018). Acetylcholine is proposed to increase the response magnitude by suppressing spontaneous activity and thereby heightening sensitivity to incoming signals (Herrero et al., 2017; Minces et al., 2017; Soma et al., 2013). In turn, noradrenaline enhances the strength and selectivity of neuronal responses to sensory input, thereby sharpening perceptual tuning curves (Kossl & Vater, 1989; McLean & Waterhouse, 1994; Waterhouse et al., 1990). Accordingly, we hypothesized that propranolol and biperiden would reduce participants’ perceptual acuity. A GLMM revealed a decreased influence of motion coherence on choices under both biperiden and propranolol, compared to placebo. While this suggests reduced perceptual abilities under both drugs, computational modeling results indicate that this effect does not reflect a genuine change in sensory processing. Instead, biperiden and propranolol seem to selectively modulate learning.

The GLMM relied on prior estimates from the Bayesian learner and therefore could not capture participants’ true trial-by-trial updating. In contrast, win-stay/lose-switch analyses allowed a model-free approximation: faster updating of the prior belief was reflected in an increased likelihood of repeating a previously rewarded choice on the next trial (win-stay), whereas previously non-rewarded choices were associated with a higher probability of switching (lose-switch). Biperiden and propranolol significantly increased the probability to repeat a previously rewarded response compared to placebo. Choices following non-rewarded responses were not significantly modulated by either drug. Computational modeling allowed us to further disentangle these effects by capturing the latent cognitive processes. Accordingly, propranolol and biperiden increased learning rates compared to placebo causing a faster updating, with no effects on the weighting of sensory evidence. Hence, prior estimates under both drugs were influenced more strongly by recent outcomes compared to more distant experiences. Conversely, we hypothesized that propranolol would decrease learning rates leading to a slower updating of prior beliefs. In a similar experimental paradigm, Lawson et al. (2021) found a slower updating of outcome contingencies under propranolol. The reason for the divergence of our results remains unclear but may be caused by differential task designs. In the task used by Lawson et al. (2021), participants did not receive explicit feedback. Instead, learning depended on the perceptual information making it difficult to fully disentangle changes in sensory processing and changes in prior updating. In contrast, in our task participants received veridical feedback following each choice, allowing learning to be based on the true outcome rather than on participants’ potentially noisy interpretation of the coherent motion. Moreover, our modeling approach explicitly estimated how perceptual information and learned beliefs were integrated to guide choices. This allowed us to dissociated the contribution of perceptual evidence from the influence of prior expectations at the decision stage.

Consistent with the observed increase in learning rates under propranolol, noradrenaline is proposed to stabilize estimates of environmental volatility (Marshall et al., 2016; Yu & Dayan, 2005). Accordingly, noradrenergic antagonism has been shown to lead to faster updating, as unexpected outcomes are more likely to be attributed to changes in the underlying contingency (Marshall et al., 2016). This is consistent with the increased learning rate observed here. However, it is important to note that prior work targeted α_1_-adrenoceptors, whereas the present study involved β-adrenoceptor antagonism. Nevertheless, it has been shown that β-adrenoceptor activity contributes to learning by supporting synaptic plasticity in the hippocampus, which is essential for memory formation (Jami et al., 2023; Larsen et al., 2023; Sara et al., 1999). We extend this finding by identifying a role of β-adrenoceptor activity in trial-by-trial learning whereby recent outcomes have a stronger influence compared to more distant experiences. Moreover, noradrenergic effects on learning have been shown to be specific to volatile environments and to track change points (Glennon et al., 2019; Jepma et al., 2016). This is consistent with our experimental paradigm, which incorporated volatility in prior probabilities, and fits the proposed noradrenergic mechanism of stabilizing estimates of the underlying contingency. Our findings further indicate that the effects of propranolol on learning were specific to rewarded outcomes, as learning rates were selectively increased following rewarded choices. Accordingly, no significant noradrenergic modulation of learning rates was observed following non-rewarded choices.

Similar to noradrenaline, cholinergic effects on learning have been shown to be specific to volatile environments (Kurtenbach et al., 2026). Recent own work has demonstrated that biperiden increases learning rates in volatile learning environments, in line with our hypothesis and current findings (Kurtenbach et al., 2026). Consistent with this, faster updating under biperiden leads prior estimates to deviate stronger from the statistical optimum compared to placebo. Hence, increased learning rates under biperiden cause maladaptive adjustments in prior beliefs. This pattern was observed both in the present study and in Kurtenbach et al. (2026). It has been proposed that muscarinic acetylcholine receptor antagonist causes impaired adaptions to probabilistic noise. More specifically, unexpected outcomes do not lead to an adequate adjustment of beliefs regarding probabilistic outcomes. This causes unexpected outcomes to be instead attributed to contingency changes, in turn causing a faster maladaptive updating (Kurtenbach et al., 2026; Marshall et al., 2016; Piray & Daw, 2021; Yu & Dayan, 2005). This interpretation is further supported by the effects of cholinergic activity in the brain. Activity of cholinergic neurons in the basal forebrain has been identified as a precise reinforcement signal (Hangya et al., 2015). Furthermore, muscarinic acetylcholine receptor antagonist by biperiden has been shown to decreases the prediction error signal correlate in high-beta power in the lateral prefrontal cortex (Kurtenbach et al., 2026). This in turn may lead to the impaired post-error adjustments that have been observed in sensory cortical areas (Danielmeier et al., 2015). Moreover, cholinergic effects on learning have been reported to be specific to rewarded choices, with only a trend-level effect for unrewarded learning rates (Kurtenbach et al., 2026). While we observe increased learning rates following both rewarded and unrewarded outcomes, differences in effects sizes, together with the win-stay/lose-switch analysis, suggest a similar pattern in our data: biperiden effects on the rewarded learning rates are larger than those on unrewarded learning rates, which only showed a small effect size. In addition, faster updating under biperiden was specific to win-stay and did not hold for lose-switch choices.

We did not find significant drug effects on the integration parameter θ, which defined the relative weighting of bottom-up sensory evidence versus top-down prior beliefs during decision making. Nevertheless, noradrenergic and cholinergic effects on learning provide insights into this integration: increased learning rates under both drugs suggests that prior beliefs are more uncertain and hence environmental contingencies need to be updated faster. Therefore, top-down prior expectations are suppressed in order to facilitate learning from bottom-up evidence, which is augmented (Yu & Dayan, 2005). This suggest that the balance between top-down prior information and bottom-up evidence is regulated indirectly by the dynamic updating of prior expectations based on the expected environmental uncertainty.

Lastly, theoretical models postulate a role of noradrenaline in guiding choice stochasticity (Doya, 2002). In our computational models, this was operationalized by an inverse temperature τ, with higher values indicating more deterministic choices. However, propranolol had no effect on choice stochasticity. Instead biperiden tended to decrease the inverse temperature, indicative of more stochastic choices. While we did not hypothesize this modulation, a similar effect of biperiden on choice stochasticity has been observed in previous work (Erfanian Abdoust et al., 2024; Kurtenbach et al., 2026).

Taken together, our findings provide causal evidence that acetylcholine and noradrenaline shape learning by modulating, in distinct ways, how prior expectations are updated in response to new information. Importantly, once differences in learning were accounted for, neither biperiden nor propranolol altered the relative weighting of sensory evidence and prior expectations during choices. These findings therefore suggest that acetylcholine and noradrenaline contribute to choice behavior by shaping learning of the internal contingencies of the environment, rather than by directly modulating the integration of sensory evidence and prior knowledge at the time of decision making.

## Methods

The study was approved by the medical ethics board of Heinrich Heine University Düsseldorf (study-ID: 2024-286) and is in line with the Helsinki Declaration of 1975. The described methods follow the planned procedure, which can be found at https://doi.org/10.17605/OSF.IO/AQD2R (Dias Maile et al., 2025). Analyses which diverged from this, were marked accordingly.

### Participants

Participants were native-level German speakers with normal or corrected-to-normal vision. They were naïve to the purpose of the study and gave written informed consent. Compensation followed a fixed rate plus an additional performance-based bonus. Due to the pharmacological intervention, participants were screened for certain pre-existing medical conditions, including neurological and psychiatric disorders (Supplement 5). As part of this screening, they were also required to reach a predefined performance threshold in the two experimental tasks. Participants who failed to meet this performance threshold across the three pharmacological sessions were excluded post hoc (*N* = 6). Of the 72 participants who took part in the study, four dropped out and six were excluded based on task performance resulting in a final sample of 62 participants (36 female). On average, they were *M* = 23.77 years old (*SD* = 3.88 years) and weight *M* = 70.28 kg (*SD* = 7.31 kg). Most participants were right-handed (*N* = 52).

### Procedure

The study consisted of a medical screening followed by three pharmacological sessions. In the screening, participants were required to meet the medical inclusion criteria (see supplements). This included an evaluation of participants’ heart rate and ECG conducted by a cardiologist to assess potential abnormalities (particularly atrioventricular block). In addition, participants filled out the Beck’s Depression Inventory (BDI; Beck et al., 1996), the State-Trait Anxiety Inventory (STAI; Spielberger et al., 1983), the Barratt Impulsiveness scale (BIS-15; Meule et al., 2011; Spinella, 2007) and a modified version of the Edinburgh Handedness Inventory (Oldfield, 1971). They also completed the two experimental tasks and needed to meet pre-defined performance criteria: at least 65% correct responses in the random dot motion task (RDM) and fewer than ten missed trials in the explore-exploit-task (EET; not further analyzed in this study).

The three pharmacological sessions were conducted in the morning and were separated by at least five days to account for the drugs’ half-lives (Borgstrom et al., 1981; Brocks, 1999; Grimaldi et al., 1986; Kalam et al., 2020). Each session lasted approximately three hours and followed an identical procedure, only the drug (or placebo) administered differed (Figure 1a). In a double-blind, within-subject, crossover design participants received a single oral dose of either the M_1_-preferring muscarinic receptor antagonist biperiden (4mg), the β-adrenoceptor antagonist propranolol (40mg), or a placebo. Prior to the drug administration, participants were provided a standardized breakfast. Vital signs (blood pressure and heart rate) were measured at three time points per session: before drug administration, immediately before the experimental tasks (45 min after drug administration), and after the experimental tasks (approximately 125 min after drug administration). At the same three time points, participants also filled out the Bond and Lader Visual Analogue Mood Scale (BL-VAS; Bond & Lader, 1974) and the STAI. Thirty minutes after drug administration, participants refamiliarized themselves with the experimental tasks. After 50 min they performed the Trail Making Test (TMT; part A; Rodewald et al., 2012). Participants began the experimental tasks one hour after drug intake in accordance with the average time for peak plasma concentration (Borgstrom et al., 1981; Brocks, 1999; Grimaldi et al., 1986; Kalam et al., 2020). Each task lasted approximately 25-35 min. Task order was counterbalanced across participants but remained constant within participant. At the end of the pharmacological session, participants had the opportunity to see the study physician. Following the third session, they were asked to indicate the perceived order of drug administration.

### Random dot motion task

In the RDM, participants viewed a random dot kinematogram in which most dots moved randomly, while a subset moved coherently either leftward or rightward (Figure 1b,c; similar to Froböse & Ort, 2022; Marshall et al., 2024). Participants were instructed to indicate the overall movement direction (left or right) with a button press.

Two aspects of the task were manipulated. First, motion coherence i.e., the proportion of coherently moving dots, was varied across four levels (6%, 12%, 24% and 48%). Second, a prior probability for motion direction was introduced. Specifically, across a given phase (44 or 84 trials), there was a probability *p* that the direction of movement would be to the right, where *p* took one of three values (0.2, 0.5, 0.8). The prior probability and its switches were not explicitly cued and therefore needed to be learned by participants. To assess performance based solely on the learned prior, four trials with 0% motion coherence were included in each prior phase. Coherence and prior were counterbalanced. The task consisted of 18 blocks with 50 trials each. Participants were allowed to rest after each block. Prior to the main task, they (re-) familiarized themselves with the task by completing two blocks featuring higher coherence levels (70%, 75%, 80% and 85%) but identical prior levels and trial structure.

In detail, each trial began with the presentation of a fixation dot (radius = 0.4 degrees visual angle (dva)), which was presented centrally for 800ms prior to the presentation of the random dot kinematogram (Thaler et al., 2013). The fixation dot remained onscreen with the kinematogram, which was presented in a ring with an inner/outer radius of 1.25/5.00 dva. On average, it consisted of 250 dots, which had a size of four pixels per dot resulting in a dot density of 3.4 dots/dva^2^. Dots moved at a speed of 3.6 dva/s and had a lifetime of four frames. Their positions were updated on every frame. After four frames, a dot was replaced by a new one appearing at a random location within the cloud. Dots were presented in light gray (RGB value: 236, 236, 236) against a dark gray background (RGB: 13, 13, 13). The random dot kinematogram was shown for 1500 ms. If no response was made during this period, another 500 ms response window followed without the stimulus on screen. Responses made after this were marked as too late. Early responses were defined as those given within 200 ms of stimulus onset. In cases of early or late responses, a warning was presented on the screen. Otherwise, feedback was provided in the form of a centrally presented smiley or frowny (radius = 0.67 dva) for 500 ms, indicating whether the response was correct. At the end of each block, participants received feedback on their percentage of correct answers. They were then given the opportunity to take a self-paced break of at least 10 s before continuing. The RDM was programmed using Psychopy (version 3.1.5; Peirce, 2007) and presented on a 16 inch screen with a refresh rate of 60 Hz. Participants were situated at a screen distance of 80 cm in a dimly lit room.

### Statistical analyses

#### General linear mixed-effects modeling

In order to evaluate drug effects on perceptual decision making and the independent weighting of perceptual evidence and prior information, behavioral data was first analyzed using GLMM. Analyses were conducted in R (version 4.2.2; R Development Core Team, 2022) using the lme4 package (version 3.1–3; Bates, Machler, et al., 2015). We chose this approach since it allowed estimating both within- and between-subject variability in accordance with our within-subject experimental design (Brown, 2021; Harrison et al., 2018; Jaeger, 2008). Using these models, we examined task and drug effects on participants’ choices and reaction times. Choice behavior (left vs. right) was modeled using a GLMM with a binomial distribution. Fixed effects included the degree of coherent motion, the learned prior and the drug condition and its interaction with both coherence and prior. Since participants did not know the true prior distribution and had to instead learn it over time, the prior was estimated individually for each participant and pharmacological condition using a Bayesian optimal learner set up in MATLAB (version R2021a; The MathWorks Inc., 2021). Deviating from the original analyses plan, we implemented the Bayesian learner described in Behrens et al. (2007) since it provided the best model fit. We applied treatment contrasts and z-scored all continuous predictors to achieve standardized estimates. The random-effects structure was determined using a principal component analysis (PCA) following the recommendation of Bates, Kliegl, et al. (2015), where random components needed to explain at least 1% variance. The choice model included both a random intercept and a random slope for the within-subject factors motion coherence, prior and drug. Exploratory analyses of learning also examined participants’ win-stay/lose-switch behavior (see supplements for a complete overview of model structure). Where we observed significant drug effects on control measures (mood, state anxiety, visuomotor abilities, heart rate and blood pressure), we added them to our GLMM as fixed effects to verify that our effects of interest were not affected by these covariates. A further control analysis encompassed adding the session number as fixed effects to control for putative order effects. All *p*-values were based on asymptotic Wald tests. *R*^2^ was reported for all models using the r.squaredGLMM procedure of the MuMIn package (version 1.48.4; Bartoń, 2024).

#### Computational modeling

To analyze the drug effects on the underlying cognitive processes (i.e., learning and the relative weighting of information sources) that determined participants’ choice behavior, we used computational modeling. Based on an independent dataset using the same perceptual decision-making task, we identified the best-fitting model. Specifically, we compared a basic model in which the influence of motion coherence and prior information was governed by a relative weighting parameter to several other implementations of this integration process. These included a model with coherence-dependent weighting, in which the influence of prior information decreased with increasing motion coherence, as well as a variant in which this dependence was exponentially scaled. We further tested a model incorporating a coherence-dependent modulation of the learning rate. Lastly, we also compared all models to a version including a side bias, given its pronounced influence on raw choice behavior. We found that the basic model with an added side bias provided the best fit to participants’ choice behavior. We therefore preregistered this model for implementation in the present pharmacological dataset. Contrary to our planned analyses, we were unable to fit the data in a Bayesian hierarchal framework due to model complexity. Therefore, the preregistered model was implemented non-hierarchically in Matlab (MATLAB Version R2021a, Massachusetts: The Mathworks Inc.). The model assumed an additive integration of the motion coherence and the learned prior:

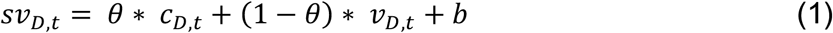

where *sv_D,t_* was the subjective value of direction *D* on trial *t*, and *c_D,t_* and *v_D,t_* represented the coherence and the learned prior (both scaled between -1 and 1), respectively. The parameter θ determined the relative weighting of these two sources of information. Values θ > 0.5 indicated that choices were more strongly influenced by the motion coherence than by the learned prior. Since GLMMs revealed a right-side bias, the model additionally included a side bias b. The learned prior was modelled using a Q-learning rule, where the current prior (*v_D,t_*) was updated for the next trial (*v_D, t+1_*) based on the outcome of the current trial (*r_D,t_*) and the learning rate λ:

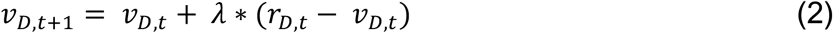

Given the specific drug effects on participants’ win-stay behavior, an exploratory model further incorporated differential learning rates based on if participants were rewarded (λ_r_) or not (λ_u_) on the current trial:

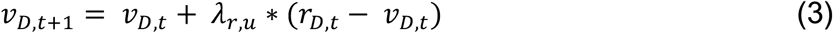

Choices were modelled using a softmax choice rule including an inverse temperature τ to capture choice stochasticity. Therefore, the probability to choose direction *D* was generated as:

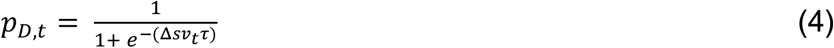

Where Δsv denoted the subjective value difference (left minus right for direction *D* = left). Because the distribution of parameter estimates deviated from normality, drug effects were assessed using Wilcoxon-signed rank tests. We additionally performed a parameter recovery and model validation to evaluate model integrity. For this purpose, we simulated 500 datasets per participant and pharmacological session. For model validation, the data was average across simulation using the mode, thereby preserving the binomial structure of the behavioral data and key behavioral findings were replicated. Parameter recovery involved refitting the model on the simulated datasets and verifying successful recovery of the ground-truth parameters.

## Code and data availability

All data, code and the preregistered analysis plan have been made publicly available on OSF (https://doi.org/10.17605/OSF.IO/AQD2R).

## Author contributions

- Study conception and design (AADM, FB, JXOR, GJ), data acquisition (AADM, EA, FB, DA, LKD), data analysis (AADM, JXOR, GJ), interpretation of data for the work (AADM, JXOR, GJ)
- Drafting the article (AADM, JXOR, GJ), revising it for intellectual content (all authors)
- Approving the final version of the manuscript (all authors)

## Acknowledgements

We thank Tomke Bartels, Hayden Bera, Johanna Dappen, Iremnaz Ercan, Julia Hoffmann, Janne Knipp, Lukas Leib, Sophie Öhlschläger and Nils Schäfer for their support during data acquisition. Computational infrastructure and support were provided by the Centre for Information and Media Technology at Heinrich Heine University Düsseldorf.

