## Supplementary Material for "Cholinergic and noradrenergic modulation of perceptual decision making and learning"

**Supplement 1: Complete mixed model fits ………..…………………………. 2**
**Supplement 2: Control analyses ……………………………………….. 5**
**Supplement 3: Complete fit for model with separate learning rates …….. 18**
**Supplement 4: Parameter recovery and model validation …………….. 19**
**Supplement 5: List of exclusion criteria ……………………………… 23**
**References ………………………………………………………………… 25**

### Supplement 1: Complete mixed model fits

Tables depicts the complete mixed model fits including a description of the fixed and random effect structure. Columns show the β-weights, SEM, z-value (t-values for linear mixed models) and p-value for the fixed effects. Linear mixed effects further include a column showing the df. Stars represent significant effects based on p < 0.05.

**Table S1 General linear mixed effects model: task effects.** Fixed effects included main effects for the intercept (side bias), z-scaled coherence and z-scaled prior (based on the Bayes optimal learner). The model included a full random-effect structure. The dependent variable was choice (right-side choice). Marginal $R^2_{GLMM} = 0.593$.

| | ***β-weight*** | ***SEM*** | ***z*-value** | ***p*-value** |
|---|---|---|---|---|
| Intercept | 0.28 | 0.03 | 8.76 | < 2e-16 * |
| Coherence | 2.08 | 0.11 | 18.42 | < 2e-16 * |
| Prior | 0.59 | 0.04 | 16.19 | < 2e-16 * |

**Table S2 Linear mixed effects model: task effects on prior.** Fixed effects included main effects for the intercept and z-scaled coherence. The model included a random intercept. The dependent variable was the difference in the probability of a correct choice between trials in which priors were aligned with versus opposed to motion coherence. Marginal $R^2_{GLMM} = 0.235$.

| | ***β-weight*** | ***SEM*** | **df** | **t-value** | ***p*-value** |
|---|---|---|---|---|---|
| Intercept | 0.40 | 0.03 | 97.82 | 17.49 | < 2e-16 * |
| Coherence | -0.66 | 0.05 | 185.00 | -13.73 | < 2e-16 * |

**Table S3 General linear mixed effects model: drug effects.** Fixed effects included main effects for the intercept (side bias), drugs, z-scaled coherence and z-scaled prior (based on the Bayes optimal learner). Additionally, fixed effects included interactions of the drugs with the z-scaled coherence and z-scaled prior. The random effects included a random intercept and slope for the main effects of z-scaled coherence, z-scaled prior and the drugs. The dependent variable was choice (right-side choice). Marginal $R^2_{GLMM} = 0.594$.

| | ***β-weight*** | ***SEM*** | ***z*-value** | ***p*-value** |
|---|---|---|---|---|
| Intercept | 0.28 | 0.04 | 7.24 | < 0.001 * |
| Biperiden | 0.03 | 0.04 | 0.69 | 0.492 |
| Propranolol | -0.03 | 0.04 | -0.71 | 0.481 |
| Coherence | 2.14 | 0.11 | 18.74 | < 2e-16 * |
| Prior | 0.57 | 0.04 | 14.83 | < 2e-16 * |
| Biperiden:coherence | -0.10 | 0.03 | -3.76 | < 0.001 * |
| Propranolol:coherence | -0.06 | 0.03 | -2.26 | 0.024 * |
| Biperiden:prior | 0.01 | 0.02 | 0.68 | 0.497 |
| Propranolol:prior | 0.06 | 0.02 | 3.23 | 0.001 * |

**Table S4 General linear mixed effects model: win-stay.** Fixed effects included main effects for the intercept (side bias), drugs, z-scaled coherence and z-scaled stay bias. Additionally, fixed effects included interactions of the drugs with the z-scaled coherence and z-scaled stay bias. The random effects included a random intercept and slope for the main effects of z-scaled coherence, z-scaled prior and the drugs. The dependent variable was choice (right-side choice). The model only included trials following a correct choice. The number of iterations was set to 100000. Marginal $R^2_{GLMM} = 0.622$.

| | ***β-weight*** | ***SEM*** | ***z*-value** | ***p*-value** |
|---|---|---|---|---|
| Intercept | 0.30 | 0.04 | 7.78 | < 0.001 * |
| Biperiden | 0.05 | 0.04 | 1.32 | 0.186 |
| Propranolol | -0.01 | 0.04 | -0.28 | 0.782 |
| Coherence | 2.55 | 0.11 | 23.03 | < 2e-16 * |
| Stay | 0.39 | 0.04 | 8.93 | < 2e-16 * |
| Biperiden:coherence | -0.12 | 0.03 | -3.72 | < 0.001 * |
| Propranolol:coherence | -0.03 | 0.03 | -0.97 | 0.333 |
| Biperiden:stay | 0.08 | 0.02 | 4.07 | < 0.001 * |
| Propranolol:stay | 0.08 | 0.02 | 4.39 | < 0.001 * |

**Table S5 General linear mixed effects model: lose-switch.** Fixed effects included main effects for the intercept (side bias), drugs, z-scaled coherence and z-scaled stay bias. Additionally, fixed effects included interactions of the drugs with the z-scaled coherence and z-scaled stay bias. Random effects included a random intercept and slope for the main effects of z-scaled coherence, z-scaled stay bias and drugs. The dependent variable was choice (right-side choice). The model only included trials following an incorrect choice. Marginal $R^2_{GLMM} = 0.568$.

| | ***β-weight*** | ***SEM*** | ***z*-value** | ***p*-value** |
|---|---|---|---|---|
| Intercept | 0.23 | 0.05 | 4.69 | < 0.001 * |
| Biperiden | -0.04 | 0.05 | -0.73 | 0.466 |
| Propranolol | -0.06 | 0.05 | -1.13 | 0.257 |
| Coherence | 2.37 | 0.11 | 20.64 | < 2e-16 * |
| Stay | -0.25 | 0.05 | -4.58 | < 0.001 * |
| Biperiden:coherence | -0.07 | 0.05 | -1.34 | 0.182 |
| Propranolol:coherence | -0.07 | 0.05 | -1.30 | 0.194 |
| Biperiden:stay | -0.04 | 0.04 | -1.02 | 0.309 |
| Propranolol:stay | 0.02 | 0.04 | 0.49 | 0.627 |

**Table S6 Linear mixed effects model: reaction times.** Fixed effects included main effects for the intercept, drugs, z-scaled absolute coherence and z-scaled congruency (coding if the Bayes optimal prior is congruent with the coherent motion). Additionally, fixed effects included interactions of the drugs with the z-scaled coherence and z-scaled congruency. Random effects included a random intercept and slope for the main effects of the drugs. The dependent variable was log-transformed reaction times. Marginal $R^2_{GLMM} = 0.077$.

| | ***β*-weight** | ***SEM*** | ***df*** | **t-value** | ***p*-value** |
|---|---|---|---|---|---|
| Intercept | -0.41 | 0.03 | 61.01 | -13.52 | < 2e-16 * |
| Biperiden | -0.01 | 0.02 | 61.00 | -0.36 | 0.721 |
| Propranolol | -0.02 | 0.02 | 61.01 | -1.16 | 0.249 |
| Coherence | -0.10 | 0.00 | 155400.00 | -80.92 | < 2e-16 * |
| Congruency | -0.02 | 0.00 | 155400.00 | -16.27 | < 2e-16 * |
| Biperiden:coherence | 0.00 | 0.00 | 155400.00 | -0.05 | 0.959 |
| Propranolol:coherence | 0.00 | 0.00 | 155400.00 | -1.00 | 0.317 |
| Biperiden:congruency | 0.00 | 0.00 | 155400.00 | 0.90 | 0.369 |
| Propranolol:congruency | 0.00 | 0.00 | 155400.00 | -0.79 | 0.431 |

### Supplement 2: Control analyses

Control analyses included capturing drug effects on participants' mood (measured with the Bond and Lader Visual Analogue Mood Scale; BL-VAS; Bond & Lader, 1974), state anxiety (measured with the State-Trait Anxiety Inventory; STAI; Spielberger et al., 1983), visuomotor abilities (measured with the Trail Making Test; TMT; Rodewald et al., 2012), blood pressure and heart rate. Mood, state anxiety and vital parameters were measured at three different time points per pharmacological session: before drug administration (baseline), 45 min after drug administration (timepoint 2) and approximately 2 h after drug administration (timepoint 3). Visuomotor abilities are tested 50 min after drug administration. Linear mixed models are implemented to test the drug effects on these measures. Significantly modulated measures are added as fixed effects to the models investigating drug effects in the experimental task to verify that our effects of interest are not affected by these covariates. Additionally, we add the session number as fixed effect to ensure that that our effects of interest are not affected by training effects. Tables depicts the complete mixed model fits including a description of the fixed and random effect structure. Columns show the β-weights, SEM, t-value (z-values for general linear mixed models) and p-value for the fixed effects. Linear mixed effects further include a column showing the df. Stars represent significant effects based on $p < 0.05$.

**Table S7 Linear mixed effects model: alertness.** Fixed effects included main effects for the intercept, measurement timepoint and drugs. Additionally, fixed effects included interactions of the drugs with the measurement timepoint. Random effects include a random intercept. The dependent variable was alertness (measured with the BL-VAS). Biperiden significantly reduces participants' alertness at measurement timepoint 3. Propranolol has no significant effect.

| | ***β-weight*** | ***SEM*** | ***df*** | ***t*-value** | ***p*-value** |
|---|---|---|---|---|---|
| Intercept | 7.84 | 0.26 | 177.30 | 29.73 | < 2e-16 * |
| Biperiden | 0.23 | 0.26 | 488.00 | 0.88 | 0.382 |
| Propranolol | -0.03 | 0.26 | 488.00 | -0.10 | 0.918 |
| Timepoint 2 | -0.27 | 0.26 | 488.00 | -1.04 | 0.298 |
| Timepoint 3 | -1.38 | 0.26 | 488.00 | -5.29 | < 0.001 * |
| Biperiden:Timepoint 2 | -0.59 | 0.37 | 488.00 | -1.60 | 0.111 |
| Propranolol:Timepoint 2 | 0.09 | 0.37 | 488.00 | 0.25 | 0.806 |
| Biperiden:Timepoint 3 | -1.52 | 0.37 | 488.00 | -4.12 | < 0.001 * |
| Propranolol:Timepoint 3 | -0.30 | 0.37 | 488.00 | -0.81 | 0.419 |

**Table S8 Linear mixed effects model: calmness.** Fixed effects included main effects for the intercept, measurement timepoint and drugs. Additionally, fixed effects included interactions of the drugs with the measurement timepoint. Random effects include a random intercept. The dependent variable was calmness (measured with the BL-VAS). The drugs have no significant effect on participants' calmness.

| | ***β-weight*** | ***SEM*** | ***df*** | ***t*-value** | ***p*-value** |
|---|---|---|---|---|---|
| Intercept | 8.34 | 0.20 | 191.35 | 41.51 | < 2e-16 * |
| Biperiden | 0.03 | 0.20 | 488.00 | 0.16 | 0.875 |
| Propranolol | 0.09 | 0.20 | 488.00 | 0.42 | 0.676 |

| | | | | | |
|---|---|---|---|---|---|
| Timepoint 2 | 0.20 | 0.20 | 488.00 | 1.00 | 0.318 |
| Timepoint 3 | 0.50 | 0.20 | 488.00 | 2.46 | 0.014 * |
| Biperiden:Timepoint 2 | 0.05 | 0.29 | 488.00 | 0.17 | 0.868 |
| Propranolol:Timepoint 2 | 0.04 | 0.29 | 488.00 | 0.09 | 0.927 |
| Biperiden:Timepoint 3 | -0.54 | 0.29 | 488.00 | -1.86 | 0.064 |
| Propranolol:Timepoint 3 | -0.13 | 0.29 | 488.00 | -0.46 | 0.647 |

**Table S9 Linear mixed effects model: contentedness.** Fixed effects included main effects for the intercept, measurement timepoint and drugs. Additionally, fixed effects included interactions of the drugs with the measurement timepoint. Random effects include a random intercept. The dependent variable was contentedness (measured with the BL-VAS). The drugs have no effect on participants' contentedness

| | *β-weight* | *SEM* | *df* | *t*-value | *p*-value |
|---|---|---|---|---|---|
| Intercept | 4.75 | 0.29 | 256.20 | 16.48 | < 2e-16 * |
| Biperiden | 0.68 | 0.32 | 488.00 | 2.11 | 0.035 * |
| Propranolol | 0.02 | 0.32 | 488.00 | 0.06 | 0.955 |
| Timepoint 2 | 0.02 | 0.32 | 488.00 | 0.06 | 0.951 |
| Timepoint 3 | -0.02 | 0.32 | 488.00 | -0.08 | 0.940 |
| Biperiden:Timepoint 2 | -0.69 | 0.45 | 488.00 | -1.52 | 0.129 |
| Propranolol:Timepoint 2 | -0.01 | 0.45 | 488.00 | -0.01 | 0.989 |
| Biperiden:Timepoint 3 | -0.78 | 0.45 | 488.00 | -1.72 | 0.085 |
| Propranolol:Timepoint 3 | 0.00 | 0.45 | 488.00 | 0.00 | 0.999 |

**Table S10 Linear mixed effects model: state anxiety.** Fixed effects included main effects for the intercept, measurement timepoint and drugs. Additionally, fixed effects included interactions of the drugs with the measurement timepoint. Random effects include a random intercept. The dependent variable was state anxiety (measured with the STAI). The drugs have no significant effect on participants' state anxiety

| | *β-weight* | *SEM* | *df* | *t*-value | *p*-value |
|---|---|---|---|---|---|
| Intercept | 29.39 | 0.78 | 110.33 | 37.71 | < 2e-16 * |
| Biperiden | 1.35 | 0.60 | 488.00 | 2.26 | 0.024 * |
| Propranolol | 0.65 | 0.60 | 488.00 | 1.08 | 0.282 |
| Timepoint 2 | -0.05 | 0.60 | 488.00 | -0.08 | 0.936 |
| Timepoint 3 | 1.03 | 0.60 | 488.00 | 1.72 | 0.085 |
| Biperiden:Timepoint 2 | -0.08 | 0.85 | 488.00 | -0.10 | 0.924 |
| Propranolol:Timepoint 2 | -0.65 | 0.85 | 488.00 | -0.76 | 0.446 |
| Biperiden:Timepoint 3 | 1.48 | 0.85 | 488.00 | 1.75 | 0.080 |
| Propranolol:Timepoint 3 | -0.47 | 0.85 | 488.00 | -0.55 | 0.581 |

**Table S11 Linear mixed effects model: visuomotor abilities.** Fixed effects included main effects for the intercept and drugs. Random effects include a random intercept. The dependent variable was visuomotor abilities (measured with the TMT). The drugs have no significant effect on participants' visuomotor abilities.

| | ***β*-weight** | ***SEM*** | ***df*** | ***t*-value** | ***p*-value** |
|---|---|---|---|---|---|
| Intercept | 25.34 | 1.23 | 108.52 | 20.67 | < 2e-16 * |
| Biperiden | 0.87 | 1.12 | 122.00 | 0.78 | 0.436 |
| Propranolol | 0.89 | 1.12 | 122.00 | 0.79 | 0.429 |

**Table S12 Linear mixed effects model: systolic blood pressure.** Fixed effects included main effects for the intercept, measurement timepoint and drugs. Additionally, fixed effects included interactions of the drugs with the measurement timepoint. Random effects include a random intercept. The dependent variable was the systolic blood pressure. Biperiden significantly reduces participants' systolic blood pressure at measurement timepoint 2. Propranolol significantly reduces participants' systolic blood pressure at measurement timepoint 2 and 3.

| | ***β*-weight** | ***SEM*** | ***df*** | ***t*-value** | ***p*-value** |
|---|---|---|---|---|---|
| Intercept | 98.29 | 2.83 | 70.72 | 34.70 | < 2e-16 * |
| Biperiden | 0.10 | 1.14 | 487.00 | 0.09 | 0.932 |
| Propranolol | 1.65 | 1.14 | 487.00 | 1.45 | 0.149 |
| Timepoint 2 | -1.23 | 1.14 | 487.00 | -1.08 | 0.282 |
| Timepoint 3 | -0.99 | 1.14 | 487.01 | -0.87 | 0.386 |
| Biperiden:Timepoint 2 | -3.31 | 1.61 | 487.00 | -2.06 | 0.040 * |
| Propranolol:Timepoint 2 | -3.35 | 1.61 | 487.00 | -2.09 | 0.038 * |
| Biperiden:Timepoint 3 | -2.25 | 1.61 | 487.01 | -1.40 | 0.163 |
| Propranolol:Timepoint 3 | -8.69 | 1.61 | 487.01 | -5.39 | < 0.001 * |

**Table S13 Linear mixed effects model: diastolic blood pressure.** Fixed effects included main effects for the intercept, measurement timepoint and drugs. Additionally, fixed effects included interactions of the drugs with the measurement timepoint. Random effects include a random intercept. The dependent variable was the diastolic blood pressure. Biperiden significantly reduces participants' diastolic blood pressure at measurement timepoint 2. Propranolol significantly reduces participants' diastolic blood pressure at measurement timepoint 3.

| | ***β*-weight** | ***SEM*** | ***df*** | ***t*-value** | ***p*-value** |
|---|---|---|---|---|---|
| Intercept | 91.26 | 2.84 | 69.83 | 32.11 | < 2e-16 * |
| Biperiden | 1.40 | 1.09 | 487.00 | 1.28 | 0.200 |
| Propranolol | 1.45 | 1.09 | 487.00 | 1.33 | 0.185 |
| Timepoint 2 | -1.63 | 1.09 | 487.00 | -1.49 | 0.137 |
| Timepoint 3 | -2.67 | 1.10 | 487.01 | -2.43 | 0.015 * |
| Biperiden:Timepoint 2 | -4.15 | 1.55 | 487.00 | -2.68 | 0.008 * |
| Propranolol:Timepoint 2 | -2.86 | 1.55 | 487.00 | -1.85 | 0.065 |
| Biperiden:Timepoint 3 | -1.75 | 1.55 | 487.01 | -1.13 | 0.259 |

| | | | | | |
|---|---|---|---|---|---|
| Propranolol:Timepoint 3 | -6.25 | 1.55 | 487.01 | -4.04 | < 0.001 * |

**Table S14 Linear mixed effects model: heart rate.** Fixed effects included main effects for the intercept, measurement timepoint and drugs. Additionally, fixed effects included interactions of the drugs with the measurement timepoint. Random effects include a random intercept. The dependent variable was the heart rate. Biperiden significantly reduces participants' heart rate at measurement timepoint 2 and 3. Propranolol significantly reduces participants' heart rate at measurement timepoint 3.

| | ***β*-weight** | ***SEM*** | ***df*** | ***t*-value** | ***p*-value** |
|---|---|---|---|---|---|
| Intercept | 74.60 | 1.25 | 146.09 | 59.80 | < 2e-16 * |
| Biperiden | 2.13 | 1.14 | 486.72 | 1.88 | 0.061 |
| Propranolol | 0.81 | 1.14 | 486.72 | 0.71 | 0.478 |
| Timepoint 2 | 0.21 | 1.14 | 486.72 | 0.19 | 0.854 |
| Timepoint 3 | -3.37 | 1.14 | 486.80 | -2.95 | 0.003 * |
| Biperiden:Timepoint 2 | -8.23 | 1.61 | 486.72 | -5.12 | < 0.001 * |
| Propranolol:Timepoint 2 | -5.47 | 1.61 | 486.72 | -3.40 | 0.001 |
| Biperiden:Timepoint 3 | -10.99 | 1.61 | 486.76 | -6.83 | < 0.001 * |
| Propranolol:Timepoint 3 | -9.91 | 1.61 | 486.76 | -6.15 | < 0.001 * |

**Table S15 General linear mixed effects model: drug effects controlling for session order.** Fixed effects included main effects for the intercept (side bias), drugs, z-scaled coherence and z-scaled prior (based on the Bayes optimal learner). Additionally, fixed effects included interactions of the drugs and z-scaled session order with the z-scaled coherence and z-scaled prior. The random effects included a random intercept and slope for the main effects of z-scaled coherence, z-scaled prior and the drugs. The dependent variable was choice (right-side choice). Drug effects in our experimental task are not affected by the session order.

| | ***β*-weight** | ***SEM*** | ***z*-value** | ***p*-value** |
|---|---|---|---|---|
| Intercept | 0.28 | 0.04 | 7.23 | < 0.001 * |
| Biperiden | 0.03 | 0.04 | 0.71 | 0.480 |
| Propranolol | -0.03 | 0.04 | -0.71 | 0.480 |
| Coherence | 2.14 | 0.11 | 18.74 | < 2e-16 * |
| Prior | 0.57 | 0.04 | 14.81 | < 2e-16 * |
| Biperiden:coherence | -0.11 | 0.03 | -3.88 | < 0.001 * |
| Propranolol:coherence | -0.07 | 0.03 | -2.34 | 0.019 * |
| Biperiden:prior | 0.02 | 0.02 | 0.81 | 0.418 |
| Propranolol:prior | 0.06 | 0.02 | 3.32 | < 0.001 * |
| Session:coherence | -0.03 | 0.01 | -2.76 | 0.006 * |
| Session:prior | 0.06 | 0.01 | 7.89 | < 0.001 * |

**Table S16 General linear mixed effects model: drug effects controlling for alertness.** Fixed effects included main effects for the intercept (side bias), drugs, z-scaled coherence, z-scaled prior (based on the Bayes optimal learner) and z-scaled alertness (difference score timepoint 1 – timepoint 3). Additionally, fixed effects included interactions of the drugs and the z-scaled coherence, z-scaled prior and z-scaled alertness. The random effects included a random intercept and slope for the main effects of z-scaled coherence, z-scaled prior and the drugs. The dependent variable was choice (right-side choice). Drug effects in our experimental task are not affected by drug effects on participants' alertness.

| | ***β-weight*** | ***SEM*** | ***z*-value** | ***p*-value** |
|---|---|---|---|---|
| Intercept | 0.27 | 0.04 | 6.60 | < 0.001 * |
| Biperiden | 0.04 | 0.04 | 1.08 | 0.279 |
| Propranolol | -0.01 | 0.04 | -0.37 | 0.712 |
| Coherence | 2.14 | 0.11 | 18.74 | < 2e-16 * |
| Prior | 0.57 | 0.04 | 14.83 | < 2e-16 * |
| Alertness | -0.05 | 0.05 | -0.92 | 0.360 |
| Biperiden:coherence | -0.10 | 0.03 | -3.76 | < 0.001 * |
| Propranolol:coherence | -0.06 | 0.03 | -2.26 | 0.024 * |
| Biperiden:prior | 0.01 | 0.02 | 0.69 | 0.493 |
| Propranolol:prior | 0.06 | 0.02 | 3.23 | 0.001 * |
| Biperiden:alertness | 0.03 | 0.06 | 0.56 | 0.576 |
| Propranolol:alertness | 0.05 | 0.06 | 0.84 | 0.401 |

**Table S17 General linear mixed effects model: drug effects controlling for systolic blood pressure at timepoint 2.** Fixed effects included main effects for the intercept (side bias), drugs, z-scaled coherence, z-scaled prior (based on the Bayes optimal learner) and z-scaled systolic blood pressure (difference score timepoint 1 – timepoint 2). Additionally, fixed effects included interactions of the drugs with the z-scaled coherence, z-scaled prior and z-scaled systolic blood pressure. The random effects included a random intercept and slope for the main effects of z-scaled coherence, z-scaled prior and the drugs. The dependent variable was choice (right-side choice). Drug effects in our experimental task are not affected by drug effects on participants' systolic blood pressure at timepoint 2.

| | ***β-weight*** | ***SEM*** | ***z*-value** | ***p*-value** |
|---|---|---|---|---|
| Intercept | 0.27 | 0.04 | 6.96 | < 0.001 * |
| Biperiden | 0.04 | 0.04 | 1.03 | 0.303 |
| Propranolol | -0.01 | 0.04 | -0.31 | 0.754 |
| Coherence | 2.14 | 0.11 | 18.77 | < 2e-16 * |
| Prior | 0.57 | 0.04 | 14.85 | < 2e-16 * |
| systole | -0.04 | 0.03 | -1.37 | 0.171 |
| Biperiden:coherence | -0.10 | 0.03 | -3.76 | < 0.001 * |
| Propranolol:coherence | -0.06 | 0.03 | -2.26 | 0.024 * |
| Biperiden:prior | 0.01 | 0.02 | 0.68 | 0.498 |
| Propranolol:prior | 0.06 | 0.02 | 3.22 | 0.001 * |

| | | | | |
|---|---|---|---|---|
| Biperiden:systole | 0.02 | 0.04 | 0.55 | 0.585 |
| Propranolol:systole | 0.02 | 0.05 | 0.35 | 0.723 |

**Table S18 General linear mixed effects model: drug effects controlling for systolic blood pressure at timepoint 3.** Fixed effects included main effects for the intercept (side bias), drugs, z-scaled coherence, z-scaled prior (based on the Bayes optimal learner) and z-scaled systolic blood pressure (difference score timepoint 1 – timepoint 3). Additionally, fixed effects included interactions of the drugs with the z-scaled coherence, z-scaled prior and z-scaled systolic blood pressure. The random effects included a random intercept and slope for the main effects of z-scaled coherence, z-scaled prior and the drugs. The dependent variable was choice (right-side choice). Drug effects in our experimental task are not affected by drug effects on participants' systolic blood pressure at timepoint 3.

| | ***β*-weight** | ***SEM*** | ***z*-value** | ***p*-value** |
|---|---|---|---|---|
| Intercept | 0.26 | 0.04 | 6.32 | < 0.001 * |
| Biperiden | 0.05 | 0.04 | 1.30 | 0.195 |
| Propranolol | 0.00 | 0.04 | 0.07 | 0.947 |
| Coherence | 2.14 | 0.11 | 18.71 | < 2e-16 * |
| Prior | 0.57 | 0.04 | 14.77 | < 2e-16 * |
| systole | -0.03 | 0.04 | -0.83 | 0.406 |
| Biperiden:coherence | -0.10 | 0.03 | -3.73 | < 0.001 * |
| Propranolol:coherence | -0.06 | 0.03 | -2.25 | 0.025 * |
| Biperiden:prior | 0.01 | 0.02 | 0.71 | 0.481 |
| Propranolol:prior | 0.06 | 0.02 | 3.25 | 0.001 * |
| Biperiden:systole | 0.06 | 0.04 | 1.36 | 0.174 |
| Propranolol:systole | 0.01 | 0.05 | 0.25 | 0.804 |

**Table S19 General linear mixed effects model: drug effects controlling for diastolic blood pressure at timepoint 2.** Fixed effects included main effects for the intercept (side bias), drugs, z-scaled coherence, z-scaled prior (based on the Bayes optimal learner) and z-scaled diastolic blood pressure (difference score timepoint 1 – timepoint 2). Additionally, fixed effects included interactions of the drugs with the z-scaled coherence, z-scaled prior and z-scaled diastolic blood pressure. The random effects included a random intercept and slope for the main effects of z-scaled coherence, z-scaled prior and the drugs. The dependent variable was choice (right-side choice). Drug effects in our experimental task are not affected by drug effects on participants' diastolic blood pressure at timepoint 2.

| | ***β*-weight** | ***SEM*** | ***z*-value** | ***p*-value** |
|---|---|---|---|---|
| Intercept | 0.27 | 0.04 | 6.66 | < 0.001 * |
| Biperiden | 0.05 | 0.04 | 1.21 | 0.225 |
| Propranolol | -0.01 | 0.04 | -0.26 | 0.798 |
| Coherence | 2.14 | 0.11 | 18.75 | < 2e-16 * |
| Prior | 0.57 | 0.04 | 14.84 | < 2e-16 * |
| diastole | -0.05 | 0.04 | -1.29 | 0.199 |
| Biperiden:coherence | -0.10 | 0.03 | -3.76 | < 0.001 * |

| | | | | |
|---|---|---|---|---|
| Propranolol:coherence | -0.06 | 0.03 | -2.27 | 0.024 * |
| Biperiden:prior | 0.01 | 0.02 | 0.68 | 0.496 |
| Propranolol:prior | 0.06 | 0.02 | 3.22 | 0.001 * |
| Biperiden:diastole | 0.02 | 0.04 | 0.36 | 0.721 |
| Propranolol:diastole | 0.01 | 0.04 | 0.20 | 0.845 |

**Table S20 General linear mixed effects model: drug effects controlling for diastolic blood pressure at timepoint 3.** Fixed effects included main effects for the intercept (side bias), drugs, z-scaled coherence, z-scaled prior (based on the Bayes optimal learner) and z-scaled diastolic blood pressure (difference score timepoint 1 – timepoint 3). Additionally, fixed effects included interactions of the drugs with the z-scaled coherence, z-scaled prior and z-scaled diastolic blood pressure. The random effects included a random intercept and slope for the main effects of z-scaled coherence, z-scaled prior and the drugs. The dependent variable was choice (right-side choice). Drug effects in our experimental task are not affected by drug effects on participants' diastolic blood pressure at timepoint 3.

| | ***β*-weight** | ***SEM*** | ***z*-value** | ***p*-value** |
|---|---|---|---|---|
| Intercept | 0.26 | 0.04 | 6.41 | < 0.001 * |
| Biperiden | 0.04 | 0.04 | 1.12 | 0.261 |
| Propranolol | 0.00 | 0.04 | -0.05 | 0.959 |
| Coherence | 2.14 | 0.11 | 18.74 | < 2e-16 * |
| Prior | 0.57 | 0.04 | 14.78 | < 2e-16 * |
| diastole | -0.04 | 0.04 | -1.03 | 0.302 |
| Biperiden:coherence | -0.10 | 0.03 | -3.74 | < 0.001 * |
| Propranolol:coherence | -0.06 | 0.03 | -2.25 | 0.025 * |
| Biperiden:prior | 0.01 | 0.02 | 0.70 | 0.483 |
| Propranolol:prior | 0.06 | 0.02 | 3.25 | 0.001 * |
| Biperiden:diastole | 0.00 | 0.05 | -0.05 | 0.964 |
| Propranolol:diastole | 0.04 | 0.05 | 0.73 | 0.464 |

**Table S21 General linear mixed effects model: drug effects controlling for heart rate at timepoint 2.** Fixed effects included main effects for the intercept (side bias), drugs, z-scaled coherence, z-scaled prior (based on the Bayes optimal learner) and z-scaled heart rate (difference score timepoint 1 – timepoint 2). Additionally, fixed effects included interactions of the drugs with the z-scaled coherence, z-scaled prior and z-scaled heart rate. The random effects included a random intercept and slope for the main effects of z-scaled coherence, z-scaled prior and the drugs. The dependent variable was choice (right-side choice). Drug effects in our experimental task are not affected by drug effects on participants' heart rate at timepoint 2.

| | ***β*-weight** | ***SEM*** | ***z*-value** | ***p*-value** |
|---|---|---|---|---|
| Intercept | 0.25 | 0.05 | 5.40 | < 0.001 * |
| Biperiden | 0.07 | 0.05 | 1.51 | 0.131 |
| Propranolol | 0.01 | 0.05 | 0.24 | 0.810 |
| Coherence | 2.14 | 0.11 | 18.78 | < 2e-16 * |

| | | | | |
|---|---|---|---|---|
| Prior | 0.57 | 0.04 | 14.84 | < 2e-16 * |
| diastole | -0.07 | 0.04 | -1.59 | 0.112 |
| Biperiden:coherence | -0.10 | 0.03 | -3.76 | < 0.001 * |
| Propranolol:coherence | -0.06 | 0.03 | -2.27 | 0.023 * |
| Biperiden:prior | 0.01 | 0.02 | 0.68 | 0.496 |
| Propranolol:prior | 0.06 | 0.02 | 3.23 | 0.001 * |
| Biperiden:heart rate | 0.06 | 0.05 | 1.11 | 0.266 |
| Propranolol:heart rate | 0.07 | 0.05 | 1.27 | 0.204 |

**Table S22 General linear mixed effects model: drug effects controlling for heart rate at timepoint 3.** Fixed effects included main effects for the intercept (side bias), drugs, z-scaled coherence, z-scaled prior (based on the Bayes optimal learner) and z-scaled heart rate (difference score timepoint 1 – timepoint 3). Additionally, fixed effects included interactions of the drugs with the z-scaled coherence, z-scaled prior and z-scaled heart rate. The random effects included a random intercept and slope for the main effects of z-scaled coherence, z-scaled prior and the drugs. The dependent variable was choice (right-side choice). Drug effects in our experimental task are not affected by drug effects on participants' heart rate at timepoint 3.

| | ***β-weight*** | ***SEM*** | ***z*-value** | ***p*-value** |
|---|---|---|---|---|
| Intercept | 0.21 | 0.05 | 4.12 | < 0.001 * |
| Biperiden | 0.10 | 0.05 | 1.88 | 0.060 |
| Propranolol | 0.05 | 0.05 | 0.99 | 0.325 |
| Coherence | 2.14 | 0.11 | 18.72 | < 2e-16 * |
| Prior | 0.57 | 0.04 | 14.77 | < 2e-16 * |
| diastole | -0.09 | 0.05 | -1.92 | 0.055 |
| Biperiden:coherence | -0.10 | 0.03 | -3.74 | < 0.001 * |
| Propranolol:coherence | -0.06 | 0.03 | -2.25 | 0.024 * |
| Biperiden:prior | 0.01 | 0.02 | 0.71 | 0.478 |
| Propranolol:prior | 0.06 | 0.02 | 3.25 | 0.001 * |
| Biperiden:heart rate | 0.10 | 0.06 | 1.64 | 0.100 |
| Propranolol:heart rate | 0.08 | 0.05 | 1.45 | 0.146 |

**Table S23 General linear mixed effects model: win-stay controlling for session order.** Fixed effects included main effects for the intercept (side bias), drugs, z-scaled coherence and z-scaled stay bias. Additionally, fixed effects included interactions of the drugs and z-scaled session order with the z-scaled coherence and z-scaled stay bias. The random effects included a random intercept and slope for the main effects of z-scaled coherence, z-scaled stay bias and the drugs. The dependent variable was choice (right-side choice). The model only included trials following a correct choice. Drug effects on win-stay behavior are not affected by the session order.

| | ***β-weight*** | ***SEM*** | ***z*-value** | ***p*-value** |
|---|---|---|---|---|
| Intercept | 0.30 | 0.04 | 7.76 | < 0.001 * |

| | | | | |
|---|---|---|---|---|
| Biperiden | 0.05 | 0.04 | 1.32 | 0.188 |
| Propranolol | -0.01 | 0.04 | -0.28 | 0.782 |
| Coherence | 2.55 | 0.11 | 23.08 | < 2e-16 * |
| Stay | 0.39 | 0.04 | 8.90 | < 2e-16 * |
| Biperiden:coherence | -0.12 | 0.03 | -3.88 | < 0.001 * |
| Propranolol:coherence | -0.03 | 0.03 | -1.07 | 0.286 |
| Biperiden:stay | 0.08 | 0.02 | 4.26 | < 0.001 * |
| Propranolol:stay | 0.08 | 0.02 | 4.38 | < 0.001 * |
| Session:coherence | -0.01 | 0.01 | -1.27 | 0.206 |
| Session:stay | 0.06 | 0.01 | 8.08 | < 0.001 * |

**Table S24 General linear mixed effects model: win-stay controlling for alertness.** Fixed effects included main effects for the intercept (side bias), drugs, z-scaled coherence, z-scaled stay bias and z-scaled alertness (difference score timepoint 1 – timepoint 3). Additionally, fixed effects included interactions of the drugs with the z-scaled coherence, z-scaled stay bias and z-scaled alertness. The random effects included a random intercept and slope for the main effects of z-scaled coherence, z-scaled prior and the drugs. The dependent variable was choice (right-side choice). The model only included trials following a correct choice. Drug effects on win-stay behavior are not affected by drug effects on participants' alertness.

| | ***β-weight*** | ***SEM*** | ***z*-value** | ***p*-value** |
|---|---|---|---|---|
| Intercept | 0.28 | 0.04 | 7.09 | < 0.001 * |
| Biperiden | 0.07 | 0.04 | 1.86 | 0.062 |
| Propranolol | 0.00 | 0.04 | 0.09 | 0.925 |
| Coherence | 2.55 | 0.11 | 23.05 | < 2e-16 * |
| Stay | 0.39 | 0.04 | 8.95 | < 2e-16 * |
| Alertness | -0.06 | 0.05 | -1.31 | 0.192 |
| Biperiden:coherence | -0.12 | 0.03 | -3.72 | < 0.001 * |
| Propranolol:coherence | -0.03 | 0.03 | -0.97 | 0.333 |
| Biperiden:stay | 0.08 | 0.02 | 4.08 | < 0.001 * |
| Propranolol:stay | 0.08 | 0.02 | 4.30 | < 0.001 * |
| Biperiden:alertness | 0.04 | 0.05 | 0.79 | 0.430 |
| Propranolol:alertness | 0.06 | 0.06 | 0.95 | 0.345 |

**Table S25 General linear mixed effects model: win-stay controlling for systolic blood pressure at timepoint 2.** Fixed effects included main effects for the intercept (side bias), drugs, z-scaled coherence, z-scaled stay bias and z-scaled systolic blood pressure (difference score timepoint 1 – timepoint 2). Additionally, fixed effects included interactions of the drugs with the z-scaled coherence, z-scaled stay bias and z-scaled systolic blood pressure. The random effects included a random intercept and slope for the main effects of z-scaled coherence, z-scaled stay bias and the drugs. The dependent variable was choice (right-side choice). The model only included trials following a correct choice. Drug effects on win-stay behavior are not affected by drug effects on participants' systolic blood pressure at timepoint 2.

| | *β-weight* | *SEM* | *z*-value | *p*-value |
|---|---|---|---|---|
| Intercept | 0.29 | 0.04 | 7.50 | < 0.001 * |
| Biperiden | 0.06 | 0.04 | 1.59 | 0.111 |
| Propranolol | 0.00 | 0.04 | 0.12 | 0.908 |
| Coherence | 2.55 | 0.11 | 22.96 | < 2e-16 * |
| Stay | 0.39 | 0.04 | 8.92 | < 2e-16 * |
| Systole | -0.04 | 0.03 | -1.23 | 0.217 |
| Biperiden:coherence | -0.12 | 0.03 | -3.73 | < 0.001 * |
| Propranolol:coherence | -0.03 | 0.03 | -0.97 | 0.333 |
| Biperiden:stay | 0.08 | 0.02 | 4.07 | < 0.001 * |
| Propranolol:stay | 0.08 | 0.02 | 4.29 | < 0.001 * |
| Biperiden:systole | 0.02 | 0.04 | 0.61 | 0.544 |
| Propranolol:systole | 0.00 | 0.04 | -0.05 | 0.964 |

**Table S26 General linear mixed effects model: win-stay controlling for systolic blood pressure at timepoint 3.** Fixed effects included main effects for the intercept (side bias), drugs, z-scaled coherence, z-scaled stay bias and z-scaled systolic blood pressure (difference score timepoint 1 – timepoint 3). Additionally, fixed effects included interactions of the drugs with the z-scaled coherence, z-scaled stay bias and systolic blood pressure. The random effects included a random intercept and slope for the main effects of z-scaled coherence, z-scaled stay bias and the drugs. The dependent variable was choice (right-side choice). The model only included trials following a correct choice. Drug effects on win-stay behavior are not affected by drug effects on participants' systolic blood pressure at timepoint 3.

| | *β-weight* | *SEM* | *z*-value | *p*-value |
|---|---|---|---|---|
| Intercept | 0.28 | 0.04 | 6.87 | < 0.001 * |
| Biperiden | 0.07 | 0.04 | 1.85 | 0.064 |
| Propranolol | 0.01 | 0.04 | 0.34 | 0.738 |
| Coherence | 2.54 | 0.11 | 22.91 | < 2e-16 * |
| Stay | 0.39 | 0.04 | 8.96 | < 2e-16 * |
| Systole | -0.03 | 0.04 | -0.75 | 0.452 |
| Biperiden:coherence | -0.11 | 0.03 | -3.69 | < 0.001 * |
| Propranolol:coherence | -0.03 | 0.03 | -0.95 | 0.343 |
| Biperiden:stay | 0.07 | 0.02 | 3.98 | < 0.001 * |
| Propranolol:stay | 0.08 | 0.02 | 4.20 | < 0.001 * |
| Biperiden:systole | 0.05 | 0.04 | 1.22 | 0.223 |
| Propranolol:systole | 0.01 | 0.05 | 0.28 | 0.782 |

**Table S27 General linear mixed effects model: win-stay controlling for diastolic blood pressure at timepoint 2.** Fixed effects included main effects for the intercept (side bias), drugs, z-scaled coherence, z-scaled stay bias and z-scaled diastolic blood pressure (difference score timepoint 1 – timepoint 2). Additionally, fixed effects included interactions of the drugs with the z-scaled coherence, z-scaled stay bias and z-scaled diastolic blood pressure. The random effects included a random intercept and slope for the main effects of z-scaled coherence, z-scaled stay bias and the drugs. The dependent variable was choice (right-side choice). The model only included trials following a correct choice. Drug effects on win-stay behavior are not affected by drug effects on participants' diastolic blood pressure at timepoint 2.

| | ***β-weight*** | ***SEM*** | ***z*-value** | ***p*-value** |
|---|---|---|---|---|
| Intercept | 0.28 | 0.04 | 7.19 | < 0.001 * |
| Biperiden | 0.07 | 0.04 | 1.79 | 0.073 |
| Propranolol | 0.01 | 0.04 | 0.18 | 0.861 |
| Coherence | 2.55 | 0.11 | 23.13 | < 2e-16 * |
| Stay | 0.39 | 0.04 | 8.95 | < 2e-16 * |
| Diastole | -0.05 | 0.04 | -1.31 | 0.190 |
| Biperiden:coherence | -0.12 | 0.03 | -3.73 | < 0.001 * |
| Propranolol:coherence | -0.03 | 0.03 | -0.97 | 0.331 |
| Biperiden:stay | 0.08 | 0.02 | 4.07 | < 0.001 * |
| Propranolol:stay | 0.08 | 0.02 | 4.29 | < 0.001 * |
| Biperiden:diastole | 0.02 | 0.04 | 0.49 | 0.625 |
| Propranolol:diastole | 0.00 | 0.04 | -0.07 | 0.945 |

**Table S28 General linear mixed effects model: win-stay controlling for diastolic blood pressure at timepoint 3.** Fixed effects included main effects for the intercept (side bias), drugs, z-scaled coherence, z-scaled stay bias and z-scaled diastolic blood pressure (difference score timepoint 1 – timepoint 3). Additionally, fixed effects included interactions of the drugs with the z-scaled coherence, z-scaled stay bias and z-scaled diastolic blood pressure. The random effects included a random intercept and slope for the main effects of z-scaled coherence, z-scaled stay bias and the drugs. The dependent variable was choice (right-side choice). The model only included trials following a correct choice. Drug effects on win-stay behavior are not affected by drug effects on participants' diastolic blood pressure at timepoint 3.

| | ***β-weight*** | ***SEM*** | ***z*-value** | ***p*-value** |
|---|---|---|---|---|
| Intercept | 0.27 | 0.04 | 6.95 | < 0.001 * |
| Biperiden | 0.07 | 0.04 | 1.82 | 0.068 |
| Propranolol | 0.01 | 0.04 | 0.30 | 0.762 |
| Coherence | 2.54 | 0.11 | 23.05 | < 2e-16 * |
| Stay | 0.39 | 0.04 | 8.98 | < 2e-16 * |
| Diastole | -0.05 | 0.04 | -1.31 | 0.189 |
| Biperiden:coherence | -0.12 | 0.03 | -3.70 | < 0.001 * |
| Propranolol:coherence | -0.03 | 0.03 | -0.95 | 0.343 |

| | | | | |
|---|---|---|---|---|
| Biperiden:stay | 0.07 | 0.02 | 3.97 | < 0.001 * |
| Propranolol:stay | 0.08 | 0.02 | 4.20 | < 0.001 * |
| Biperiden:diastole | 0.02 | 0.05 | 0.36 | 0.721 |
| Propranolol:diastole | 0.06 | 0.05 | 1.15 | 0.248 |

**Table S29 General linear mixed effects model: win-stay controlling for heart rate at timepoint 2.** Fixed effects included main effects for the intercept (side bias), drugs, z-scaled coherence, z-scaled stay bias and z-scaled heart rate (difference score timepoint 1 – timepoint 2). Additionally, fixed effects included interactions of the drugs with the z-scaled coherence, z-scaled stay bias and heart rate. The random effects included a random intercept and slope for the main effects of z-scaled coherence, z-scaled stay bias and the drugs. The dependent variable was choice (right-side choice). The model only included trials following a correct choice. Drug effects on win-stay behavior are not affected by drug effects on participants' heart rate at timepoint 2.

| | ***β*-weight** | ***SEM*** | ***z*-value** | ***p*-value** |
|---|---|---|---|---|
| Intercept | 0.26 | 0.04 | 5.89 | < 0.001 * |
| Biperiden | 0.09 | 0.04 | 1.96 | > 0.050 |
| Propranolol | 0.03 | 0.05 | 0.57 | 0.570 |
| Coherence | 2.55 | 0.11 | 22.89 | < 2e-16 * |
| Stay | 0.39 | 0.04 | 8.92 | < 2e-16 * |
| Diastole | -0.06 | 0.04 | -1.48 | 0.139 |
| Biperiden:coherence | -0.12 | 0.03 | -3.73 | < 0.001 * |
| Propranolol:coherence | -0.03 | 0.03 | -0.97 | 0.331 |
| Biperiden:stay | 0.08 | 0.02 | 4.07 | < 0.001 * |
| Propranolol:stay | 0.08 | 0.02 | 4.30 | < 0.001 * |
| Biperiden:heart rate | 0.05 | 0.05 | 1.07 | 0.285 |
| Propranolol:heart rate | 0.05 | 0.05 | 0.92 | 0.355 |

**Table S30 General linear mixed effects model: win-stay controlling for heart rate at timepoint 3.** Fixed effects included main effects for the intercept (side bias), drugs, z-scaled coherence, z-scaled stay bias and z-scaled heart rate (difference score timepoint 1 – timepoint 3). Additionally, fixed effects included interactions of the drugs with the z-scaled coherence, z-scaled stay bias and z-scaled heart rate. The random effects included a random intercept and slope for the main effects of z-scaled coherence, z-scaled stay bias and the drugs. The dependent variable was choice (right-side choice). The model only included trials following a correct choice. Drug effects on win-stay behavior are not affected by drug effects on participants' heart rate at timepoint 3.

| | ***β*-weight** | ***SEM*** | ***z*-value** | ***p*-value** |
|---|---|---|---|---|
| Intercept | 0.23 | 0.05 | 4.54 | < 0.001 * |
| Biperiden | 0.11 | 0.05 | 2.24 | 0.025 * |
| Propranolol | 0.06 | 0.05 | 1.28 | 0.200 |
| Coherence | 2.54 | 0.11 | 23.05 | < 2e-16 * |
| Stay | 0.39 | 0.04 | 8.97 | < 2e-16 * |

| | | | | |
|---|---|---|---|---|
| Diastole | -0.09 | 0.05 | -1.92 | 0.055 |
| Biperiden:coherence | -0.12 | 0.03 | -3.71 | < 0.001 * |
| Propranolol:coherence | -0.03 | 0.03 | -0.95 | 0.340 |
| Biperiden:stay | 0.07 | 0.02 | 3.97 | < 0.001 * |
| Propranolol:stay | 0.08 | 0.02 | 4.20 | < 0.001 * |
| Biperiden:heart rate | 0.11 | 0.06 | 1.84 | 0.066 |
| Propranolol:heart rate | 0.08 | 0.05 | 1.44 | 0.149 |

### Supplement 3: Complete model fits of computational model with separate learning rates

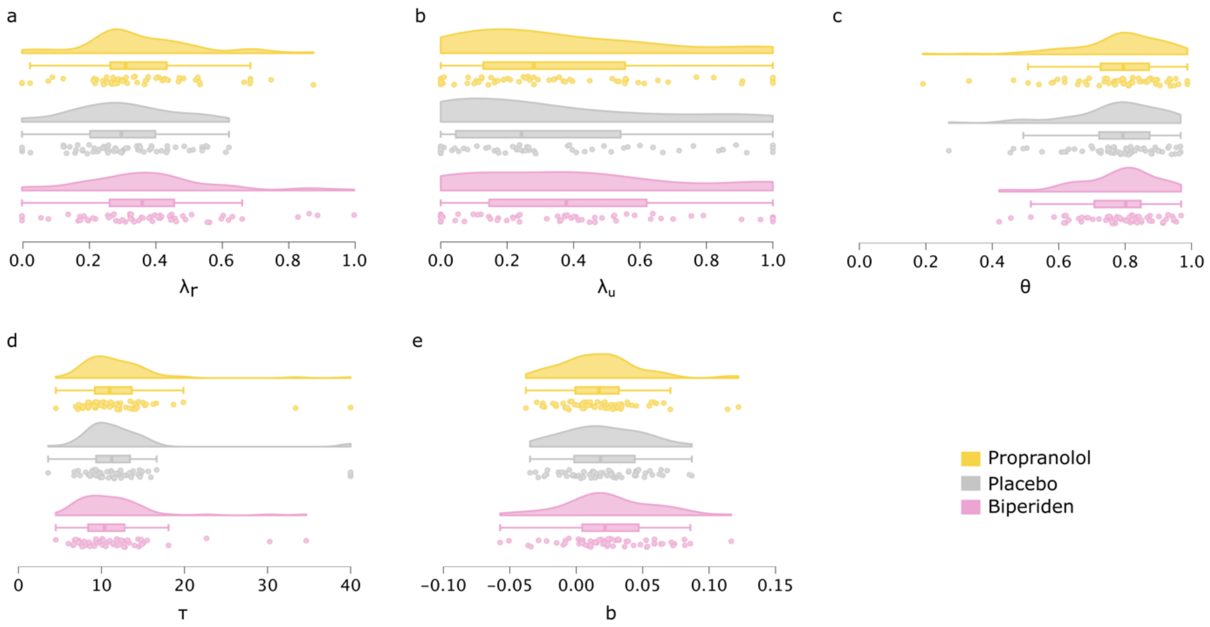


**Figure S1: Model fit of computational model with separate learning rate for rewarded and unrewarded trials.** Raincloud plots in **a**, **d**, **c**, **d**, **e** illustrate the distribution of the five model parameters across drug conditions: **(a)** learning rate for rewarded trials ($\lambda_r$), **(d)** learning rate for unrewarded trials ($\lambda_u$), **(c)** relative weighting ($\theta$), **(d)** inverse softmax temperature ($\tau$) and **(e)** right-side bias (b).

### Supplement 4: Model validation and parameter recovery

We performed a model validation and parameter recovery for both computational models (one learning rate vs. two separate learning rates) to ensure the reliability of the fitting procedure. Based on the parameter fits of the computational models, 500 dataset per participant and pharmacological session were simulated. Model validation included reproducing key behavioral findings based on the simulated data. Parameter recovery entailed fitting the computational models on the simulated data and recovering the ground-truth parameters.

#### Best fitting Model with one learning rate

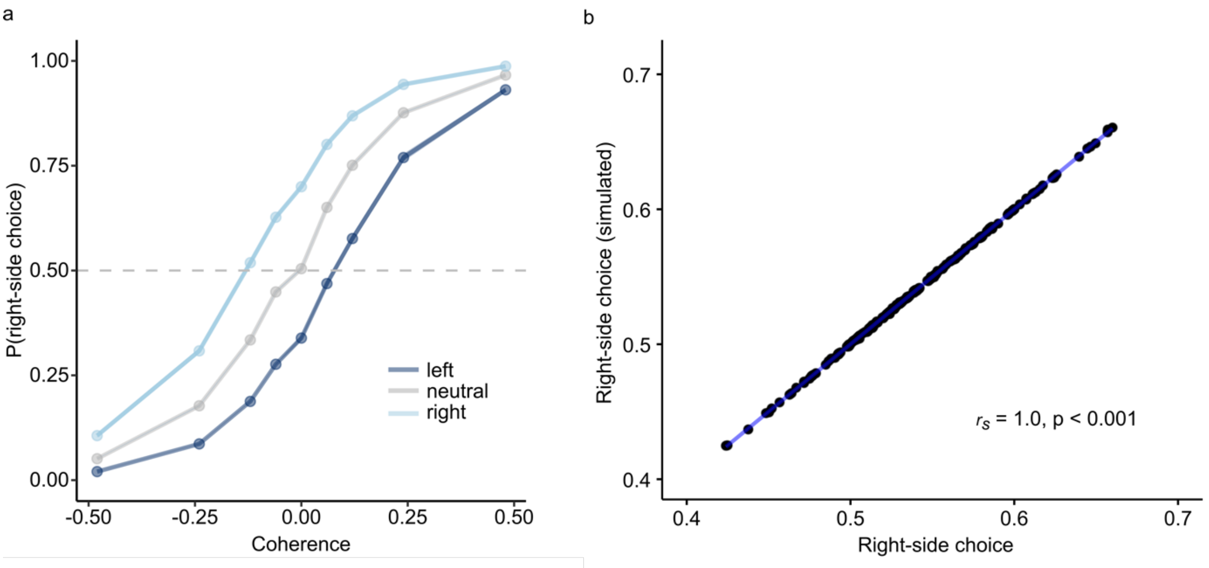


**Figure S2: Model validation.** Simulated data can reproduce the behavioral pattern. **(a)** simulated behavior average across pharmacological condition. Plotted is the probability for a right-side choice as a function of motion coherence (x-axis) and prior belief (shades of blue). Negative coherence levels correspond to leftward motion. As right-side motion increases, participants are more likely to choose the right side. Further, participants show a higher probability of right-side choice when expecting right-side motion (right prior). Shades around the graphs represent the standard error. **(b)** Spearman correlation between the probability of a right-side choice for simulated and participants' data.

**Table S31 General linear mixed effects model: simulated data.** The model was run on the simulated data whereby the mode was used to average across simulations (to keep the binary data structure). Fixed effects included main effects for the intercept (side bias), drugs, z-scaled coherence and z-scaled prior (based on the Bayes optimal learner). Additionally, fixed effects included interactions of the drugs with the z-scaled coherence and z-scaled prior. The random effects included a random intercept and slope for the main effects of z-scaled coherence, z-scaled prior and the drugs. The dependent variable was choice (right-side choice). In order to enable model convergence with the same random effects structure as in the original model (TableS2), we needed to use the "bobyqa" optimizer. However, results did not differentiate compared to the default optimizer. We can successfully reproduce the task effects as well as the negative interaction of both drugs with coherence.

| | ***β-weight*** | ***SEM*** | ***z*-value** | ***p*-value** |
|---|---|---|---|---|
| Intercept | 1.29 | 0.29 | 4.49 | < 0.001 * |
| Biperiden | 0.42 | 0.24 | 1.73 | 0.084 |
| Propranolol | -0.06 | 0.23 | -0.27 | 0.786 |
| Coherence | 12.67 | 1.25 | 10.10 | < 2e-16 * |
| Prior | 3.20 | 0.14 | 23.59 | < 2e-16 * |

| | | | | |
|---|---|---|---|---|
| Biperiden:coherence | -0.36 | 0.10 | -3.68 | < 0.001 * |
| Propranolol:coherence | -0.84 | 0.09 | -9.18 | < 2e-16 * |
| Biperiden:prior | -0.06 | 0.06 | -0.97 | 0.333 |
| Propranolol:prior | -0.05 | 0.06 | -0.80 | 0.425 |

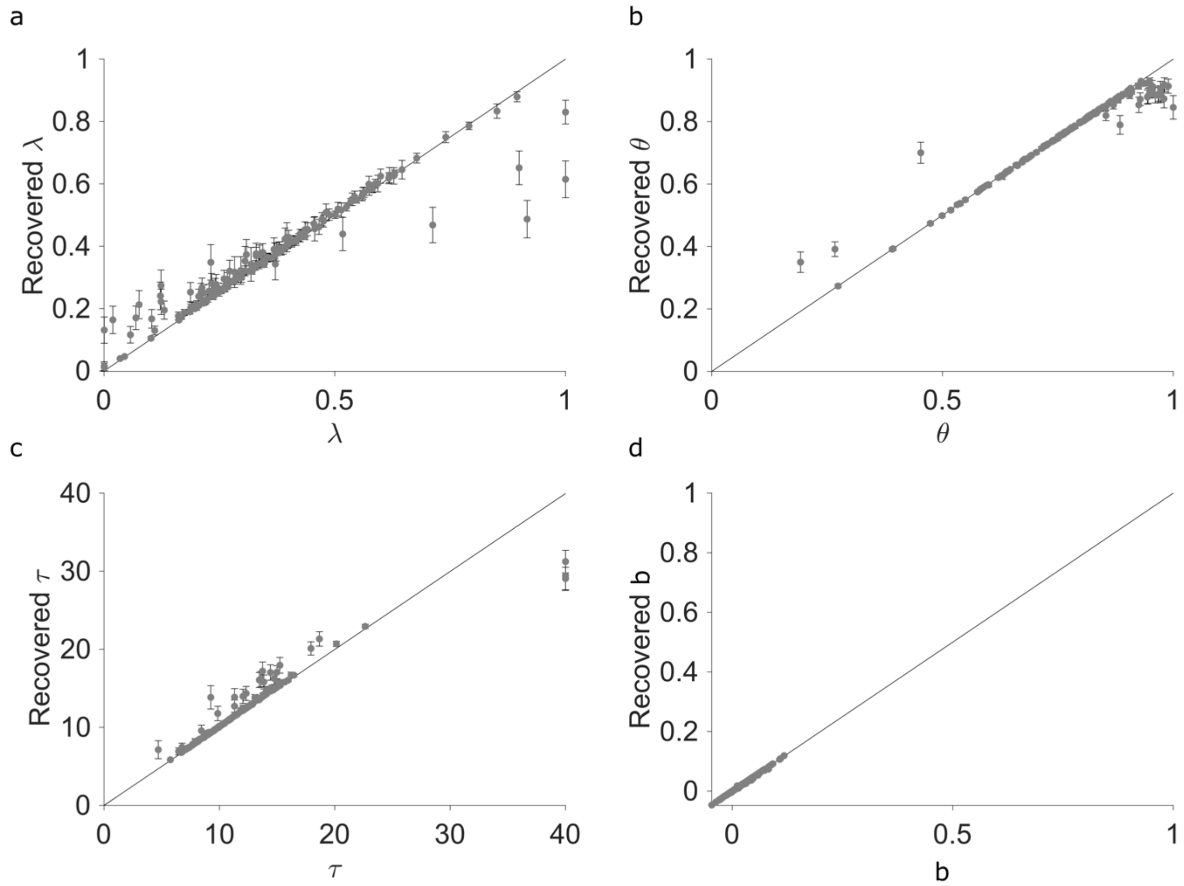


**Figure S3: Parameter recovery.** Present are the ground-truth parameter fits (x-axis) as a function of the recovered parameter fits (y-axis) for the four model parameters: $\lambda$ **(a)**, $\theta$ **(b)**, $\tau$ **(c)**, b **(d)**. Parameter fits based on the simulated data can recover the ground truth parameters.

### Model with separate learning rates

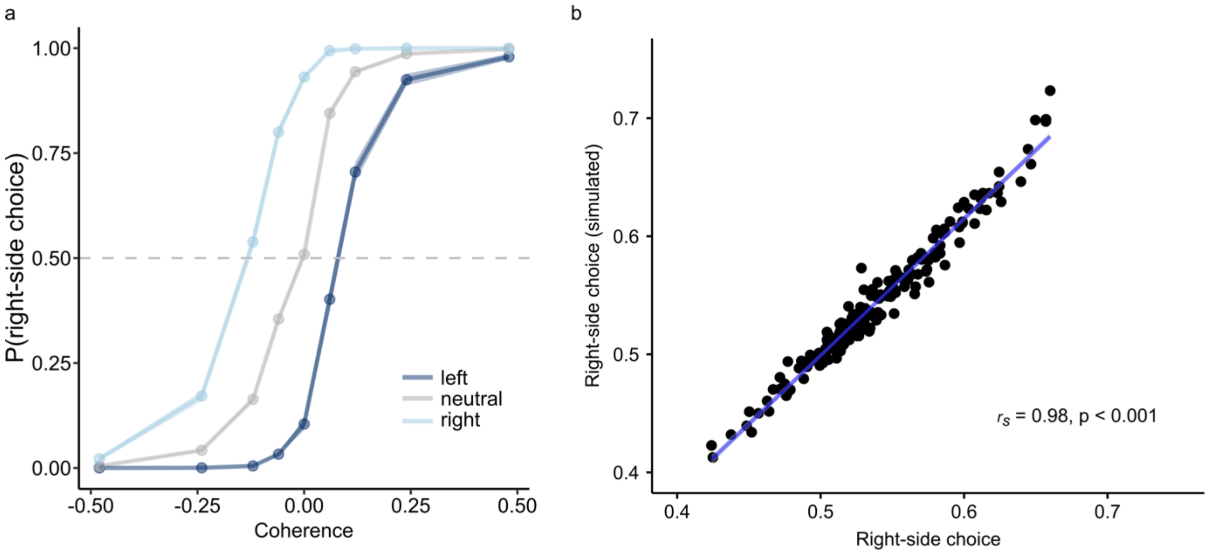


**Figure S4: Model validation.** Simulated data can reproduce the behavioral pattern. **(a)** simulated behavior average across pharmacological condition. Plotted is the probability for a right-side choice as a function of motion coherence (x-axis) and prior belief (shades of blue). Negative coherence levels correspond to leftward motion. As right-side motion increases, participants are more likely to choose the right side. Further, participants show a higher probability of right-side choice when expecting right-side motion (right prior). Shades around the graphs represent the standard error. **(b)** Spearman correlation between the probability of a right-side choice for simulated and participants' data.

**Table S32 General linear mixed effects model: simulated data.** The model was run on the simulated data whereby the mode was used to average across simulations (to keep the binary data structure). Fixed effects included main effects for the intercept (side bias), drugs, z-scaled coherence and z-scaled prior (based on the Bayes optimal learner). Additionally, fixed effects included interactions of the drugs with the z-scaled coherence and z-scaled prior. The random effects included a random intercept and slope for the main effects of z-scaled coherence, z-scaled prior and the drugs. The dependent variable was choice (right-side choice). In order to enable model convergence with the same random effects structure as in the original model (TableS2), we needed to use the "bobyqa" optimizer. However, results did not differentiate compared to the default optimizer. We can successfully reproduce key behavioral findings including the negative interaction of both drugs with coherence.

| | ***β*-weight** | ***SEM*** | ***z*-value** | ***p*-value** |
|---|---|---|---|---|
| Intercept | 1.20 | 0.24 | 5.04 | < 0.001 * |
| Biperiden | 0.23 | 0.20 | 1.16 | 0.245 |
| Propranolol | -0.10 | 0.20 | -0.53 | 0.595 |
| Coherence | 10.70 | 0.99 | 10.82 | < 2e-16 * |
| Prior | 2.74 | 0.12 | 22.80 | < 2e-16 * |
| Biperiden:coherence | -0.20 | 0.09 | -2.34 | 0.020 * |
| Propranolol:coherence | -0.64 | 0.08 | -7.89 | < 0.001 * |
| Biperiden:prior | 0.04 | 0.06 | 0.69 | 0.491 |
| Propranolol:prior | 0.07 | 0.05 | 1.21 | 0.226 |

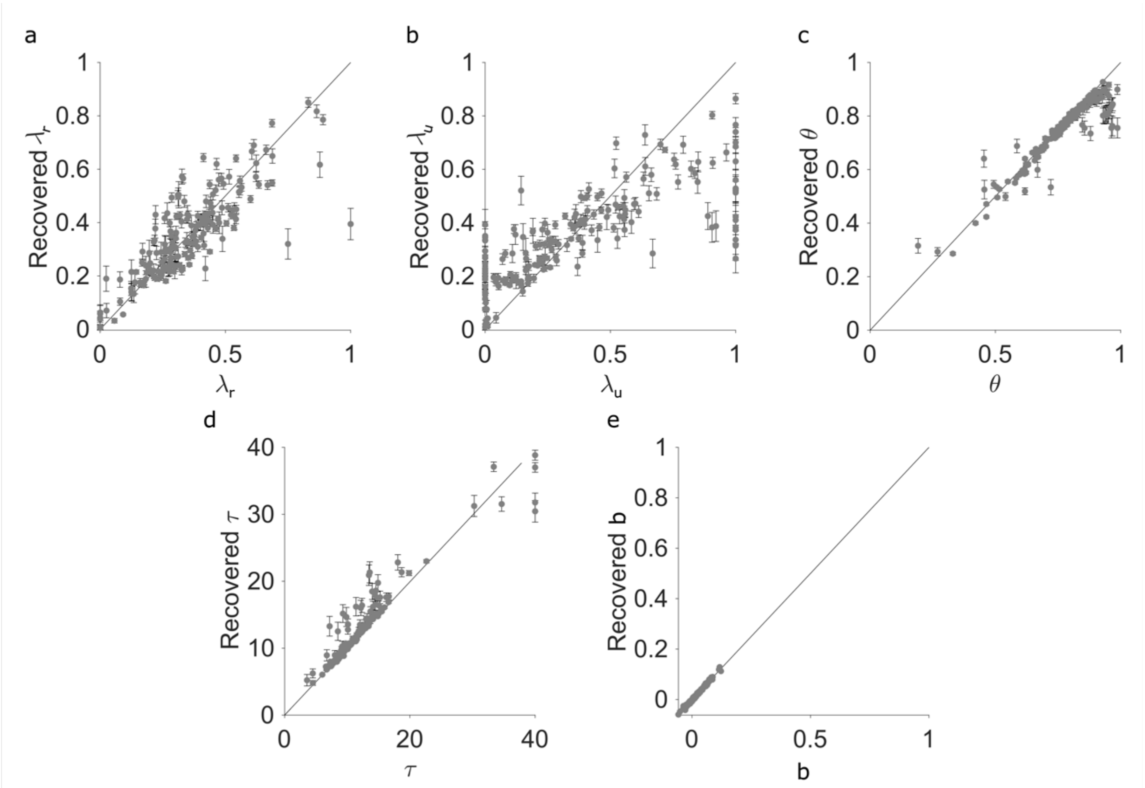


**Figure S5: Parameter recovery.** Present are the ground-truth parameter fits (x-axis) as a function of the recovered parameter fits (y-axis) for the five model parameters: $\lambda_r$ **(a)**, $\lambda_u$ **(b)**, $\theta$ **(c)**, $\tau$ **(d)**, b **(e)**. Parameter fits based on the simulated data can recover the ground truth parameters.

**Supplement 5: Exclusion criteria**

- German skills below B2 level
- Age < 18 years or > 35 years
- Weight < 50 kg or > 90
- BMI < 18.5 or > 28.0
- Lactose intolerance
- Possibility of pregnancy or current pregnancy
- Breastfeeding
- Hypersensitivity to the active ingredients or other components of biperiden, propranolol, or other beta-receptor blockers
- Heart rate < 60 bpm
- Smoking > 6 cigarettes per day
- Consumption of >13 units of alcohol (man)/ > 10 units of alcohol (women)
- Significant history of substance abuse (cannabis, amphetamines, cocaine, ecstasy, barbiturates, tranquilizers, opiates, psychedelics, etc.)
    - "Significant" is defined as
        - Consumption of any of the above substances within the last month
        - More than occasional consumption (> 5 times in a lifetime) of any of the above substances with the exception of cannabis
        - Regular (more than once a month on average) use of cannabis
- Significant heart conditions in (grand)parents and siblings (e.g., heart attack or bradycardic cardiac arrhythmia before the age of 70)
- Cardiovascular diseases (hypertension, hypotension, cardiac arrhythmias, hemophilia, heart failure, heart block, acidosis, Late stages of peripheral circulatory disorders, pulmonary obstructions, conditions that can lead to tachycardia, etc.)
- Current or previous treatment for psychiatric or neurological disorders
- Unexplained loss of consciousness in the past
- Head injury in the past
- Epilepsy
- Diabetes
- Asthma
- Bronchial spasms
- Thyroid dysfunction
- Glaucoma or angle-closure glaucoma
- Ileus
- Megacolon
- Stenosis of gastrointestinal tract
- Stomach or duodenal ulcer
- (History of) Bleeding in the gastrointestinal tract
- Organ dysfunction (e.g., renal dysfunction or liver failure)
- Prostate adenoma with residual urine formation
- Shock
- Performance-based exclusion:

- RDM: performance < 65% both in the screening and across the three pharmacological sessions
- EET: > 10 missed trials in the screening
